# Strain-level antigen variation facilitates immune evasion in *Bacteroides thetaiotaomicron*

**DOI:** 10.1101/2024.12.20.629425

**Authors:** Robert W.P. Glowacki, Jessica M. Till, Orion D. Brock, Vladimir Makarov, Morgan J. Engelhart, Philip P. Ahern

## Abstract

The T cell receptor (TCR) repertoire of intestinal CD4+ T cells is enriched for specificity towards microbiome-encoded epitopes shared among many microbiome members, providing broad microbial reactivity from a limited pool of cells. These cells actively coordinate mutualistic host-microbiome interactions, yet many epitopes are shared between gut symbionts and closely related pathobionts and pathogens. Given the disparate impacts of these agents on host health, intestinal CD4+ T cells must maintain strain-level discriminatory power to ensure protective immunity while preventing inappropriate responses against symbionts. However, to date, the mechanisms by which this occurs have remained enigmatic. To interrogate this, we leveraged BθOM mice that express a transgenic TCR specific for a *BT4295*-encoded epitope in *B. thetaiotaomicron*. While many *B. thetaiotaomicron* strains potently activated BθOM CD4+ T cells *in vitro*, strain dnLKV9 escaped recognition. Bioinformatic analyses uncovered two *BT4295* homologs in *B. thetaiotaomicron*-dnLKV9, with each homolog harboring sequence modifications relative to strain VPI-5482, specifically a premature stop codon and a T548S substitution within the epitope. Reconstruction of these variants in *B. thetaiotaomicron*-VPI-5482^Δ*BT4295*^ conferred evasion from BθOM CD4+ T cells *in vitro* to this otherwise permissive strain. Adoptive transfer of BθOM CD4+ T cells to gnotobiotic RAG1−/− colonized with *B. thetaiotaomicron* harboring these variant *BT4295* forms verified the sufficiency of these antigen modifications for evasion of BθOM CD4+ T cells. Collectively, these data uncover the existence of strain-level immune evasion in *B. thetaiotaomicron* and reveal a mechanism whereby strains evade recognition by CD4+ T cells, facilitating strain-level discrimination in responsiveness to the microbiome.

## Introduction

The intestine is home to a complex microbial ecosystem, the gut microbiome, that contributes to organismal health [1–3]. The network of intestinal immune cells is continuously exposed to microbial products to which they must remain tolerant. CD4+ T cells play an integral role in maintaining intestinal homeostasis through their capacity to adopt specialized functions that both regulate inflammatory responses and limit the penetration of microbial products into the lamina propria [1, 2, 4–8]. As such, a variety of microbiome members can coordinate the development of intestinal Th1, Th2, Th17 [9–12], regulatory (Treg) [13–18] or follicular helper (Tfh) T cells [19, 20] that ensure the maintenance of mutualistic interactions with the microbiome. Although these microbes can promote environments that are conducive to differentiation of CD4+ T cell subsets irrespective of whether the cells are specific for antigens from the inciting microbe [21–23], several studies have demonstrated that the CD4+ T cell populations induced by these microbiome members express T cell receptors (TCR) that are specific to antigens derived from these agents [19, 20, 24–37]. For example, Segmented Filamentous Bacteria (SFB) is a potent inducer of Th17 cells in the small intestinal lamina propria, and the Th17 cells triggered by SFB harbor TCRs that are specific for SFB antigens [9, 10, 24, 27]. Similarly, several species of *Helicobacter* can drive the development of Treg responses in the colonic lamina propria, and this is associated with the emergence of *Helicobacter*-specific T cells within the Treg pool [25, 26, 30]. Thus, specific cellular fates are tied to TCR specificity for members of the microbiome, thus linking the functions of a given T cell subset to the presence of individual microbes.

However, this presents challenges for intestinal CD4+ T cells. The large number of potential epitopes that can be derived from the gut microbiome is enormous and although there need not be a T cell clone specific for every epitope, the extraordinary epitope diversity coupled to constraints on the size of the CD4+ T cell compartment, dictate a need for an efficient system that balances the requirement for reactivity to a broad array of microbes with a limited number of cells. The concept of immunodominance of particular epitopes [38] and the inherent cross-reactivity of the TCR [39] represent mechanisms through which the constrained size of the intestinal T cell compartment could possess reactivity to multiple microbiome members (i.e. not all theoretical epitopes will be responded to, and one cell could recognize multiple epitopes). However, early studies showing that microbiome-derived immunodominant epitopes could be shared among phylogenetically related microbes [28, 29] provided an additional potential explanation for how a limited set of CD4+ T cells could retain activity to a broad microbial community through recognition of many microbes via a single TCR-epitope pair. Indeed, a systematic assessment of the specificities among intestinal T cells showed that the TCR pool in intestinal CD4+ T cells is dominated by TCRs that are specific for epitopes that are widely shared and broadly distributed across multiple members of the microbiome [40]. Collectively, these studies provide a conceptual framework to understand how the intestinal CD4+ T cell compartment can retain reactivity to a broad array of microbes despite the constraints on the number of T cell clones that can be maintained.

Despite the utility of this model, it does not adequately account for the need to tailor the quality of the CD4+ T cell response in a highly specific manner, often at the level of the individual strain from which the epitope is derived. For example, different strains of gut microbiome species will have substantial genomic similarity yet may impart distinct effects on the host, with some mediating beneficial functions while others have deleterious effects, exemplified by toxigenic and non-toxigenic strains of *Bacteroides fragilis* [41–45]. Indeed, there is potentially significant antigen overlap between gut symbionts and overt pathogens as is evidenced by the existence of gut-resident commensal strains of *Escherichia coli* and related enteropathogenic *E. coli* and *Shigella* [46, 47]. Thus, intestinal CD4+ T cells must maintain the capacity to deploy their inflammatory function in response to some microbes and tolerogenic properties in response to others, even if two such agents share the same epitope. However, how such discriminatory power is achieved has remained unclear. Indeed, whether or not all strains that genomically encode an epitope elicit responsiveness of T cell clones specific for the epitope, and promote the same quality of response, is largely unknown.

One reason for these limitations has been a lack of systems to adequately interrogate these questions. Although some groups have used heterologous expression of defined T cell epitopes in different bacteria to determine the impact on intestinal T cell responses [27, 48], the dearth of information regarding naturally existing shared epitopes in genetically tractable gut microbes has limited progress in this area. Here we have leveraged the BθOM TCR transgenic mouse line which expresses an I-A^b^-restricted TCR specific for a defined epitope present in the gut symbiont *Bacteroides thetaiotaomicron* (*B. theta* hereafter) [32]. The combination of transgenic TCR clone and the array of strains of *B. theta* with sequenced genomes, coupled with its genetic tractability facilitate the high-resolution required to define antigen variation across strains and how it impacts detection by CD4+ T cells. Using this system, we found that genomic encoding of the epitope recognized by BθOM CD4+ T cells by different *B. theta* isolates was not sufficient for T cell activation, and that strain *B. theta*-dnLKV9 lacked the capacity to stimulate BθOM CD4+ T cells at levels above that of a *B. theta* strain deficient in the epitope, despite expression of the gene at levels equivalent to strains that were potent stimulators of BθOM CD4+ T cells. Using a CD4+ T cell transfer system in gnotobiotic RAG1−/− mice, we found that this strain also failed to provide stimulation to these cells *in vivo* validating the evasion of BθOM CD4+ T cells by *B. theta*-dnLKV9. Bioinformatic interrogation of the genome of *B. theta*-dnLKV9 uncovered: (i) the presence of a premature stop codon downstream of the epitope recognized by BθOM CD4+ T cells, and (ii) the existence of an additional homolog of *BT4295*, present at a distinct locus in the genome that harbored a substitution, T548S, that severely impacted the capacity of the epitope to stimulate BθOM CD4+ T cells. Reconstruction of these two forms of the antigen in a strain background that supports stimulation of BθOM CD4+ T cells, but where endogenous *BT4295* had been deleted, confirmed that these sequence variations were sufficient to confer the BθOM CD4+ T cell evading properties of *B. theta*-dnLKV9. These data uncover the existence and mechanistic basis through which select bacterial strains may evade CD4+ T cell responses specific for epitopes they encode. Thus, while these data suggest that individual T cell clones may recognize many gut bacterial strains, the constellation of clones that are specific for a strain will be unique, thus providing a means through which species-specific clones and strain-specific clones allow recognition of a broad array of microbes while retaining strain-level resolution in the ensuing responses as needed.

## Materials and Methods

### Mice

C57BL/6J (Jackson Labs, strain # 000664) were purchased from Jackson Labs as needed and allowed to acclimatize in a non-barrier facility for at least 1 week prior to use. BθOM mice were provided to us by Paul Allen (Washington University in St. Louis) and maintained in-house following rederivation in a non-barrier facility. BθOM mice were on a RAG1−/−CD45.1+/+CD45.2−/− background. Conventional mice (non-gnotobiotic) were housed in a non-barrier facility under a 14 hour light and 10 hour dark cycle, and provided with food (LabDiet, Cat. # 5010 – Laboratory Autoclavable Rodent Diet) and water *ad libitum*. Breeder mice were fed irradiated Teklad global 19% protein extruded diet (Teklad Laboratory Animal Diets, Cat. # 2019) and at weaning mice were switched to irradiated 2018 Teklad Global 18% Protein Rodent Diet (Teklad Laboratory Animal Diets, Cat. # 2018). Cages contained 1/8 inch corncob bedding and nestlets were provided for enrichment. The following agents are tolerated in this facility: Mouse Norovirus (MNV), Murine Chapparvoviruses (MuCPV/MKPV), *Helicobacter* spp., *Corynebacterium bovis*, *Rodentibacter* spp., *Entamoeba muris*, and *Tritrichomonas* spp. RAG1−/− (B6.129S7-Rag1tm1Mom/J; Jackson Labs, strain # 002216) were purchased from Jackson Labs and rederived as germ-free in-house, and a germ-free colony was maintained by intercrossing RAG1−/− mice. Germ-free mice were bred in plastic flexible film isolators (Class Biologically Clean, Ltd., Madison, WI) under a 14 hour light and 10 hour dark cycle. Food (LabDiet, Cat. # 5021 for breeder mice and Cat. # 5010 for weaned mice) and water were provided *ad libitum.* For germ-free mice, all food, bedding, and water (non-acidified) were sterilized by autoclaving. Cages contained ALPHA-dri Plus Bedding (Shepherd Specialty Papers, Cat. # ALPHA-dri+PLUS), and paper towels were provided for enrichment. Sterility of germ-free mice breeding in isolators is ascertained monthly by the Gnotobiotics Facility using tests that rotate between aerobic and anaerobic cultures on rich media, Gram stains of fecal smears, and 16S rRNA PCR. In addition, mold traps are also maintained in each isolator. Mice were housed in Allentown Sentry Sealed Positive Pressure cages (Allentown) which were held in an Allentown Housing Unit for execution of experiments. All experiments and procedures were approved by the Institutional Animal Care and Use Committee (IACUC) of the Cleveland Clinic Foundation. Both male and female mice were used for experiments shown. During the course of this work, we uncovered that BθOM mice in our colony harbored a deletion in *Il10*. Whole genome sequencing identified a variety of other mutations relative to the reference C57BL/6J genome. These mutations, and the associated methods and details are described in **Table S1A-S1D**.

### Media formulations for murine cells

All assays involving the culture of murine cells were performed in “complete” Roswell Park Memorial Institute Medium (RPMI) that was formulated as follows: RPMI 1640 supplemented with HEPES, Fetal Bovine Serum, penicillin/streptomycin, and 2-mercaptoethanol as indicated in **Table S1E**. Media was sterilized by passing through a 0.22 micron filter (Fisher Scientific, Cat. # FB12566510) to sterilize prior to use. For the isolation of cells from the colon, “digestion RPMI” was prepared in the same manner as complete RPMI but 2-mercaptoethanol was not added and the media was not filter-sterilized.

### Bacterial growth and genetic manipulation of *B. theta* stains

*B. thetaiotaomicron* strains were routinely grown in tryptone-yeast extract-glucose (TYG) media (see **Table 1 F-1G** for components) that was sterile filtered prior to use using a 0.22 micron filter unit (Fisher Scientific, Cat. # FB12566510) and allowed to equilibrate in an anaerobic chamber (Coy Laboratory, Grass Lake, MI; atmosphere 10% carbon dioxide, 5% hydrogen, balance nitrogen) for 2 days prior to use. For maintenance of *B. thetaiotaomicron* on solid media, brain-heart-infusion (BHI)-blood agar plates were prepared in Petri dishes (Fisher, Cat. # FB0875712) by autoclaving BHI Broth (BD, Cat. # 211059) and agar (Fluka, Cat. # BP1423) that had been mixed with MilliQ water up to 90% of the desired end volume, followed by the addition of defibrinated horse blood (Quad Five, Cat. # 210-500) to a final concentration of 10% v/v once media had cooled to ∼60°C after autoclaving. Where necessary for genetic manipulation antibiotics and selective compounds were added to BHI-blood plates as follows: 200 µg/mL gentamicin (Gibco, Cat. # 15750060), 25 µg/mL erythromycin (Sigma, Cat. # E5389), and 200 µg/mL 5-fluoro-2’-deoxyuridine (FUdR) (Matrix Scientific, Cat. # 0089451G). *E. coli* strains used in genetic manipulation of *B. thetaiotaomicron* were grown overnight aerobically with agitation at 200 rpm in lysogeny broth (LB) broth Millers modification (Fisher Scientific, Cat. # BP1426) with 300 µg/mL of ampicillin (Sigma, Cat. # A9518) added. In-frame genetic manipulation of *B. thetaiotaomicron* strain VPI-5482 were done via allelic exchange through homologous recombination using pExchange as described previously [49]. Briefly, all manipulations in VPI-5482 were done in a strain lacking thymidine kinase (Δ*tdk*) as well as*BT4295* (Δ*BT4295*) [32] to facilitate counter-selection using the toxic drug, FUdR (102) and to insert different constructs of *BT4295* with the desired modifications to the epitope. Briefly, S17 *E. coli* λpir harboring the plasmid/construct of interest and *B. thetaiotaomicron* strains were grown separately overnight and then subcultured the following day and grown to an OD_600_ of ∼0.6-0.8 representing a mid-log phase of growth. At this point the *E. coli* and *B. thetaiotaomicron* were combined, centrifuged at 8,000 x g for 5 minutes and resuspended in 1 mL of PBS. This resuspension was then plated on BHI-blood plates with no antibiotic added and incubated at 37°C aerobically to allow for the conjugative transfer of the plasmid from *E. coli* into *B. thetaiotaomicron*. After ∼16 hr of incubation, the resulting dense lawn of bacteria was scraped from the plate and resuspended in PBS before being plated on BHI-blood plates containing gentamicin to select for only *B. thetaiotaomicron* species (the *E. coli* strain used is susceptible to gentamicin) and erythromycin to select for positive *B. thetaiotaomicron* transformants harboring the appropriate plasmid, plates were incubated anaerobically for three days. Following this, individual colonies were picked into TYG without antibiotics and grown overnight in liquid media (these represent the merodiploid stage) these were then plated on BHI-blood plates containing 200 ug/mL FUdR as a counter-selection as those cells that have not lost the plasmid (which encodes a copy of the thymidine kinase gene) will be unable to grow at this stage due to FUdR toxicity in the presence of a functional thymidine kinase gene. Individual colonies were then grown in TYG and DNA isolated and screened by PCR and Sanger sequencing to confirm gene deletions. All strains, primers, and plasmids are listed in **Table S1H-1J**.

### Preparation of bacteria for *in vitro* assays

To prepare bacteria for *in vitro* assays, *B. thetaiotaomicron* strains were grown overnight in TYG media to stationary phase as described above. 1 mL was pelleted by centrifugation at 10,000 x g, supernatant removed, pellet resuspended in PBS, cells pelleted again by centrifugation at 10,000 x g and supernatant removed. Cells were then resuspended in 1 mL PBS and heat-killed by incubation at 95°C for 90 minutes. Prior to use in assays, cells were pelleted and resuspended in the appropriate volume of complete RPMI and normalized using OD_600_. For graphs showing *in vitro* activation with OD_600_ equivalent units on the x-axis, these values represent the total OD_600_ equivalent units present in the well at the indicated value, i.e. if a culture of *B. theta* had an OD_600_=4, then 50 microlitres of culture, washed and added to a well with a final end volume of 200 microlitres following addition of BθOM CD4+ T cells would represent and OD_600_ unit of 1 in a well.

### CD4+ T cell isolation

CD4+ T cells used for *in vitro* assays and *in vivo* transfer experiments were isolated from the spleen. Spleens were harvested and stored on ice in Ca2^+^-free and Mg2^+^-free phosphate buffered saline (PBS) (prepared in-house from Caisson, Cat. # PBP01) supplemented with 0.1% w/v bovine serum albumin (BSA) (Sigma, Cat. # A3059-500G) (PBS/BSA hereafter). Splenocytes were isolated as previously described [50]; briefly, spleens were physically forced through 70 micron (Falcon; Cat. # 352350) or 100 micron (Falcon; Cat. # 352360) sterile cell strainers, and pelleted by centrifugation at 453 X g for 5 minutes. The supernatant was decanted and red blood cells were lysed by addition of ∼750 microlitres pre-warmed (37C) ammonium-chloride-potassium (ACK) lysis buffer (Gibco, Cat. # A10492-01) followed by immediate mixing. After 3 minutes lysis was halted by adding ∼10 mL of PBS/BSA to each tube, and cells were filtered through a 70– or 100-micron sterile cell strainer and centrifuged at 453 X g for 5 minutes. Supernatant was poured off and pellet resuspended for downstream use. For *in vitro* BθOM CD4+ T cell isolation, naïve CD4+ T cells were enriched using the Miltenyi Mouse Naïve CD4+ T cell Purification Kit (Miltenyi Biotec, Cat. # 130-104-453) as per the manufacturer’s instructions. For *in vivo* transfer experiments, wild-type CD4+ T cells were isolated using the Miltenyi Mouse CD4+ T cell isolation kit (Miltenyi Biotec, Cat. # 130-104-454) as per the manufacturer’s instructions. For *in vivo* transfer of BθOM CD4+ T cells, splenic naïve CD4+ T cells were purified as CD4^+^TCRVβ12^+^CD45.1^+^CD25^-^CD44^lo^CD62L^+^ or CD4^+^TCRVβ12^+^CD45.1^+^CD45.2^-^CD25^-^CD44^lo^CD62L^+^ using a FACSAria Fusion cell sorter. All steps in preparation of wild-type and BθOM CD4+ T cells for transfer *in vivo* were performed in PBS/BSA supplemented with penicillin/streptomycin as per complete RPMI. Cells were purified using a FACSAria Fusion cell sorter, and sorted into complete RPMI (10% v/v fetal bovine serum). The purity of CD4+ cells ranged from ∼91-94% (see **Fig S1** for examples of pre-and post-sorting). Of the purified CD4+ T cells, ∼95%-97% were CD25^-^CD44^lo^CD62L^+^. Wild-type CD4+ T cells and BθOM CD4+ T cells were both washed twice in PBS by pelleting at 453 X g for 5 minutes, decanting of supernatant and resuspending the pellet in ∼15 mL of PBS. Cells were mixed at a ratio of ∼95% wild-type CD4+ T cells wild-type plus ∼5% BθOM CD4+ T cells in PBS. Cells were transferred via the retro-orbital route as follows: mice were anesthetized using isoflurane, a drop of proparacaine hydrochloride 0.5% (Bausch + Lomb, Cat. # NDC24208-730-06) was administered to the eye to be injected prior to injection, cells were then injected in a volume of 150 microlitres, following which an additional drop of proparacaine hydrochloride was administered. The absolute cell number transferred differed slightly between experiments based on cell yield after purification/enrichment, ranging from 3X10^4^ BθOM + 47 X10^4^ wild-type cells for study of *B. theta*-5482 versus dnLKV9 to 2.8X10^4^ BθOM + 47.2 X10^4^ wild-type cells for study of the wild-type or T548S form of BT4295. All mice in a given experiment were administered the same number of cells.

### Bone Marrow-Derived Dendritic Cell (BMDC) Preparation

To generate BMDC, the bone marrow of wild-type C57BL/6J mice was isolated, red blood cells were lysed following the same procedure used for spleens (with 7 minute centrifugation steps), using a volume of 1 mL ACK lysis buffer to lyse red blood cells. Bone marrow cells were then seeded in a polystyrene Petri dish (Fisher, Cat. # FB0875712) at a density of 2 × 10^6^ cells in 10 mL complete RPMI 1640 supplemented with 20 ng/mL granulocyte-macrophage colony-stimulating factor (GM-CSF) (PeproTech, Cat. # 315-03) on Day 0. On Day 3 an additional 10 mL of complete RPMI supplemented with 20 ng/mL GM-CSF was carefully added with gentle swirling to mix. On Day 6 and Day 9, 10 mL was withdrawn from the edge of each plate, centrifuged at 453 X g for 5 minutes, supernatant removed and the pellet resuspended in 10 mL of complete RPMI supplemented with 10 ng/mL or 5 ng/mL GM-CSF respectively, and 10 mL added back to the plate followed by gentle swirling of the Petri dish to mix. Cells were harvested for use in assays from Day 10-11 by pipetting of non-adhered/loosely adhered cells, centrifugation at 453 X g, discarding supernatant and resuspending the pellet in complete RPMI or PBS, pelleting again at 453 X g, removing supernatant and resuspending cells in complete RPMI for use in downstream assays.

### Colony forming unit (CFUs) assay

Enumeration of CFUs from bacterial cultures was performed by spot-dilution plating on BHI-blood agar plates prepared as above with addition of 200 µg/mL gentamicin (Gibco, Cat. # 15750060). 10 µL of overnight culture was normalized to an OD_600_ of 1.0 and serially diluted 1:10 and then 10 µL of these dilutions were spot plated onto plates in technical duplicate and allowed to incubate 48-72 hours anaerobically followed by visual counting of colonies. Colony forming unit (CFU) enumeration from *in vivo* transfer experiments was performed as follows: flash frozen fecal pellets were weighed in a tared 2 mL microcentrifuge tube and resuspended at 100 mg/mL in PBS followed by agitation via pipetting, vortexing, and manual disruption of the pellet using a sterile wood applicator. Particulate matter was allowed to settle ∼5-10 minutes at room temperature. 1:10 serial dilutions were performed and spot plated on BHI-blood plates and incubated for 48-96 hours. Total CFU were calculated by multiplying colony counts by the relevant dilution factor and accounting for the plated volume such that CFU/ml were obtained. Data were then normalized based on the concentration of feces/ml to obtain CFU counts/g of feces.

### Quantitative Reverse Transcriptase PCR measurements of *BT4295*

*B. thetaiotaomicron* strains were grown overnight in sterile pre-reduced TYG, subcultured 1:100 into 5 mL of fresh TYG media the next day and grown to stationary phase (OD_600_ of ∼1.2-1.6) in TYG. 1.5 mL of cells were then spun down at 8,000 x g for 3 minutes, resuspended in 500 µL ribonucleic acid (RNA) protect bacterial reagent (Qiagen, Cat. # 76506), allowed to incubate at room temperature for 10 minutes followed by centrifugation at 5,000 x g for 10 minutes. RNA protect was removed and the pellets were stored at −80°C until processed further. Total RNA was extracted using buffers from the RNeasy Mini Kit (Qiagen) and purified on RNA-binding spin columns (Epoch Life Science, Cat. # 1940-250). Purified RNA was treated with 4 units of DNase I (1µL/unit) (NEB, Cat. # M0303) for 1 hour at 37°C. After DNase inactivation using 4 µL of 0.5 M ethylenediaminetetraacetic acid (EDTA) (Thermo Fisher Scientific, Cat. #AM9260G), RNA was purified again using a second RNA-binding spin column. Reverse transcription was performed using SuperScript III reverse transcriptase (Invitrogen, Cat. # 18080093) and random primers (Invitrogen, Cat. # 48190011). The abundance of each target *BT4295* homolog transcript in the resulting cDNA was quantified by quantitative PCR using an in-house qPCR mix as previously described [51]. Briefly, each 20 µL reaction contained 1X Thermopol Reaction Buffer (NEB, Cat. # B9004S), 125 µM dNTPs (Thermo Fisher Scientific, Cat. # 10297018), 2.5 mM MgSO4 (NEB, Cat. # B1003S), 1X SYBR Green I (Lonza, Cat. # 50513), 500 nM gene specific or 65 nM 16S rRNA primer specific to *Bacteroides* species [49] and 0.5 units Hot Start Taq Polymerase (NEB, Cat. # M0495S). 10 ng of template cDNA was used per reaction. To determine gene expression levels, the expression of the 16S rRNA was used as a housekeeping gene, and fold change was calculated relative to those obtained for the type strain, VPI-5482 using the ΔΔCT method. Primers are listed in **Table S1H.**

### *In vitro* BθOM CD4+ T cell assays

To determine the capacity of various strains of *B. theta* to stimulate BθOM CD4+ T cells, BMDC were prepared as above and incubated with heat-killed *B. theta* at a variety of doses overnight at 37°C in a volume of 100 microlitres of complete RPMI in a humidified incubator supplemented with 5% C02. Incubations were performed in flat-bottomed 96 well plates (Falcon, Cat. # 353072). BθOM CD4+ T cells were isolated as described above and added in a volume of 100 microlitres with BMDC (1:1 ratio of T cells and BMDC) and mixed by pipetting. 2 days later cells were prepared for flow cytometric-based analysis. The precise number of cells seeded varied depending on yield, from ∼5X10^4^-1X10^5^ of each cell type per well. Cell numbers were identical for all groups in a given experiment. For experiments showing stimulation with synthetic peptides, the wild-type (EEFNLPTTNGGHAT) and T548S (EEFNLPTSNGGHAT) forms of the peptides were commercially synthesized (GenScript) and purified (>98% purity) and added to wells at the indicated concentrations overnight as for bacteria prior to the addition of BθOM CD4+ T cells. Independent experiments were represented by distinct sets of BMDC, BθOM CD4+ T cells and batches of heat-killed bacteria.

### Isolation of colonic cells

To isolate cells from the colon for phenotyping, colons were extracted immediately after euthanasia, opened longitudinally, and rinsed in PBS/BSA by vigorous swirling. Colons were then chopped into 4-5 pieces and stored in ∼10 mL of PBS/BSA in a 50 mL tube (Falcon, Cat. # 352070) on ice until ready for further processing. To remove epithelial cells and intraepithelial lymphocytes, the majority of the PBS/BSA solution was removed (residual buffer remained to ensure the tissue was fully immersed in liquid) and ∼15 mL of PBS supplemented with EDTA (final concentration 5 mM; Corning, Cat. # 10823013) was added (PBS/EDTA hereafter). Tubes were placed laterally in an orbital shaker and incubated at 37°C with rotation of 180 rpm for 15-20 minutes. Tubes were then vigorously shaken for ∼30 seconds, tissue was allowed to settle and the majority of the PBS/EDTA was removed. An additional 15-20 mL of PBS/EDTA was added and the incubation step repeated. Tubes were again vigorously shaken for ∼30 seconds, following which the tissue was removed, briefly dunked in PBS/BSA before being minced with razor blades. To remove residual EDTA the minced tissue was then placed in a 50 mL tube containing ∼10 mL of digestion RPMI (complete RPMI without addition of 2-mercaptoethanol), and the tube was gently inverted to mix. After 5 minutes of incubation, the majority of the digestion RPMI was removed and a fresh ∼10 mL of digestion RPMI added, tube was gently inverted to mix and then allowed to sit at room temperature for 5 additional minutes. The majority of the digestion RPMI was removed and 10 mL of digestion RPMI supplemented with 0.1 mg/mL Collagenase Type VIII (Sigma, Cat. # C2139) and 0.075 U/mL Dispase (Corning, Cat. # 345235) was added. Tubes were placed laterally in an orbital shaker and incubated at 37°C with rotation of 180 rpm for 30 minutes. Following this incubation, tubes were vortexed for ∼30 seconds, tissue allowed to settle and the majority of the digestion RPMI was harvested, and passed through a 70 micron cell strainer into a fresh 50 mL tube. The remaining tissue underwent a second identical round of digestion with digestion RPMI supplemented with collagenase type VIII and dispase. The harvested digestion media was pooled with that of the first digestion by passing through a 70 micron cell strainer, and ∼15 mL PBS/EDTA was added to halt the digestion. Cells were pelleted by centrifugation at 453 X g for 10 minutes, followed by resuspension in ∼45 mL PBS/BSA. Cells were again pelleted by centrifugation, supernatant removed, and cells resuspended in the residual PBS/BSA volume in the tube, and stored on ice until further use.

### Flow cytometry

For flow cytometric based analysis, cells were placed in round bottom plates (Falcon, Cat. # 353077) and washed by centrifugation at 453 X g for 5 minutes, supernatant removed by flicking the solution out of the plate. Cells were then resuspended by pipetting in 50 microlitres of a solution of PBS/BSA containing anti-CD16/CD32 antibodies and a fixable viability dye (see **Table S1K** for all flow cytometry reagent details), and stored at 4C for 20-30 minutes. Cells were then pelleted as described for the wash step, and resuspended by pipetting in 50 microlitres of a staining cocktail containing the relevant fluorophore-conjugated antibodies, and stored at 4C for 20-30 minutes. Cells were then washed by addition of 200 microlitres PBS/BSA on top of the cells, followed by centrifugation and supernatant removal as described above. This step was repeated two additional times prior to fixation. For fixation, cells were resuspended by pipetting in 100 microlitres of a commercially available formaldehyde-based fixative (BioLegend, Cat. # 420801; used when no intracellular staining was subsequently performed) or (ThermoFisher, Cat. # 00-5123-43 and 00-5223-56 mixed at a 1:3 ratio respectively; used when intracellular staining was subsequently performed). Cells were stored overnight at 4C, and fixative was removed by the addition of 200 microlitres PBS/BSA on top of the cells, followed by centrifugation at 652 x g and supernatant removal as described above (note, centrifuge speed was increased once cells were fixed). This step was repeated one additional time. Cells were then either resuspended in 200 microlitres PBS/BSA for acquisition or intracellular staining was performed. For intracellular staining, cells were next permeabilized by resuspending them in 50μL permeabilization buffer (1:10 dilution of permeabilization buffer [ThermoFisher, Cat. # 00-8333-56] in Milli-Q water, supplemented with 2% volume/volume normal rat serum [SIGMASIGMASIGMA]) per well and incubating for 30min in the dark at 4°C. Cells were then spun down and re-suspended in 50μL of intracellular staining mix which contained antibodies targeting intracellular targets diluted in permeabilization buffer. Cells were incubated in the intracellular staining mix for 30min in the dark at 4°C. Unbound intracellular stain was washed off by centrifuging and washing cells twice in 200μL of permeabilization buffer, with resuspension of cells by pipetting up and down during each wash step. Cells were washed a final time in PBS/BSA, centrifuged, and resuspended in 200μL of PBS/BSA and transferred to flow tubes (Sigma, Cat. #CLS4401) for sample acquisition. All samples were acquired using a LSRII Fortessa (BD Biosciences) with BD FACSDiva software. All data was analyzed in FlowJo software (versions 10.10.0).

For cell isolation by flow cytometry, staining was performed similarly to described above, with the following changes: (i) staining was performed in 50 mL tubes with cells resuspended at ∼1X10^8^ cells/mL, (ii) Fc blocking reagents were not removed prior to incubation with fluorophore-conjugated antibodies. Cells were purified by cell sorting on a FACSAria Fusion.

### Comparative genomics of BT4295 homologs

Homolog alignment for BT4295 protein was performed by using NCBI protein-protein BLAST (Basic Local Alignment Search Tool) [52] (for the full amino acid sequence of BT4295 found in strain VPI-5482). The top BLAST results of 95% or greater similarity and at least 90% coverage were included in manual curation of a list of homologs, we confined our search to isolates and excluded sequences found in metagenomic datasets. With recent large depositions of *B. theta* isolates, our search still yielded over 100 hits with significant homology. We therefore limited our alignment to a subset of 36 isolates that were either present in our BLAST search or verified through Sanger sequencing to contain BT4295 with the BθOM T-cell epitope. Sanger sequencing was done on strains for which we have isolates of, and to distinguish between BT4295 homologs found in strain dnLKV9. Full protein sequences were downloaded and aligned in MegAlign Pro v.17.1.1 (DNASTAR, Inc) using the MUSCLE algorithm [53].

### Anti-BT4295 anti-sera and ELISA

A rabbit anti-BT4295 anti-sera was generated at Cocalico Biologicals Inc. by immunization with custom-synthesized BT4295 protein containing 600 amino acids with the predicted N-terminal signal peptide removed and an N-terminal 6xHis tag added. As determined by SDS-PAGE under reducing conditions, the purity was assayed to be >90%. (GenScript) together with Freund’s Adjuvant System. Cocalico Biologicals Inc. maintains a current USDA research license (23-R-0089) and a current Animal Welfare Assurance with NIH’s Office of Laboratory Animal Welfare (D16-00398 (A3669-01)). The expression of BT4295 by wild-type and mutant forms of *B. thetaiotaomicron* was determined by ELISA as follows: High-binding polystyrene half-area 96 well plates (Corning, 3690) were coated overnight with 50 microlitres of a solution of the *B. thetaiotaomicron* strain of interest (OD_600_ = ∼2) at 4C. To facilitate quantification, standards consisting of the purified BT4295 protein used for immunization were plated as per the bacterial cells. The next day the plates were washed at least 3 times by filling wells with wash buffer comprised of Ca2+-free and Mg+-free PBS (MP Biomedicals, 2810305) supplemented with Tween® 20 to a final concentration of 0.05% v/v (Sigma-Aldrich, P9416) (wash buffer hereafter), followed by removal of wash buffer. Wells were blocked by the addition of 125 microlitres of PBS supplemented with FBS (final concentration 10% v/v) for 2 hours at room temperature with shaking. Wells were again washed at least three times, following which 50 microliters of primary anti-sera (1:10,000 dilution) was added and incubated for 2 hours at room temperature with shaking. Once again, wells were washed at least three times, followed by the addition of 50 microlitres of a horseradish peroxidase-conjugated anti-rabbit IgG detection antibody (Invitrogen, Cat. # 31460) was added at a 1:7500 dilution in PBS containing 3% FBS v/v and allowed to incubate for 90 minutes at room temperature in the dark. Wells were washed at least 5 times before the addition of 50 microlitres of TMB substrate prepared as per the manufacturer’s instructions (BioLegend, 421101). The wells were visually monitored for color change and the ELISA was stopped by the addition of 50 microlitres of H_2_SO_4_, typically ∼10-20 minutes after the addition of substrate. Absorbance was read at 450nm on a Synergy HT plate reader (BioTek). Analysis of resulting ELISA values was done in GraphPad Prism (v.10.2.1).

### Statistical Analysis

Statistical analyses were performed using GraphPad Prism (version 10.2.1.). The specific statistical tests used are indicated within the figure legends. P values <0.05 were considered to be statistically significant.

## Results

### *B. theta* strain dnLKV9 is not recognized by BθOM CD4+ T cells

To investigate whether a given TCR that recognizes a microbiome-derived antigen would provide the capacity to recognize all microbes that express the antigen, or whether particular microbes may evade such recognition we leveraged the previously established “BθOM” TCR transgenic mouse system [32]. BθOM mice express a transgene-encoded I-Ab restricted T cell receptor (TCR) comprised of TCRVα1 and TCRVβ12 chains that recognizes an epitope (EEFNLPTTNGGHAT) encoded by *BT4295* in the type strain, *B. theta*-VPI-5482. This TCR is largely specific for *B. theta* as the epitope is not broadly distributed among members of the *Bacteroides* [32], and its target is packaged into outer membrane vesicles (OMV) that mediate its delivery across the gut epithelium [31]. Using an *in vitro* system, we set out to interrogate whether the presence of *BT4295* (or homologs thereof) would confer the capacity to be detected by BθOM CD4+ T cells. We therefore leveraged a panel of *B. theta* strains that carry a copy of *BT4295* to probe this in an *in vitro* system. Bone-marrow derived dendritic cells (BMDC) were pre-loaded with heat-killed strains of *B. theta* and mixed with naïve CD4+ T cells from BθOM mice. The capacity of BθOM CD4+ T cells to recognize these isolates was then assessed by measurement of CD69 expression on T cells, an early marker of T cell activation [54, 55], after 2 days of co-culture (**Fig 1A)**. Stimulation with the type strain of *B. theta* (VPI-5482) led to a dose-dependent increase in the proportion of cells expressing CD69 as previously reported [32] (**Fig 1B and Fig S2A)**. Stimulation with additional strains of *B. theta*, strain 0940-1 or 5951, induced similar dose-dependent increases in CD69 expression, equivalent to the type strain, in keeping with the idea that BθOM CD4+ T cells would be broadly reactive to isolates of *B. theta* (**Fig 1B**). However, stimulation with strain dnLKV9 (described in [31, 56]) failed to elicit CD69 expression to an appreciable degree.

**Figure 1.**
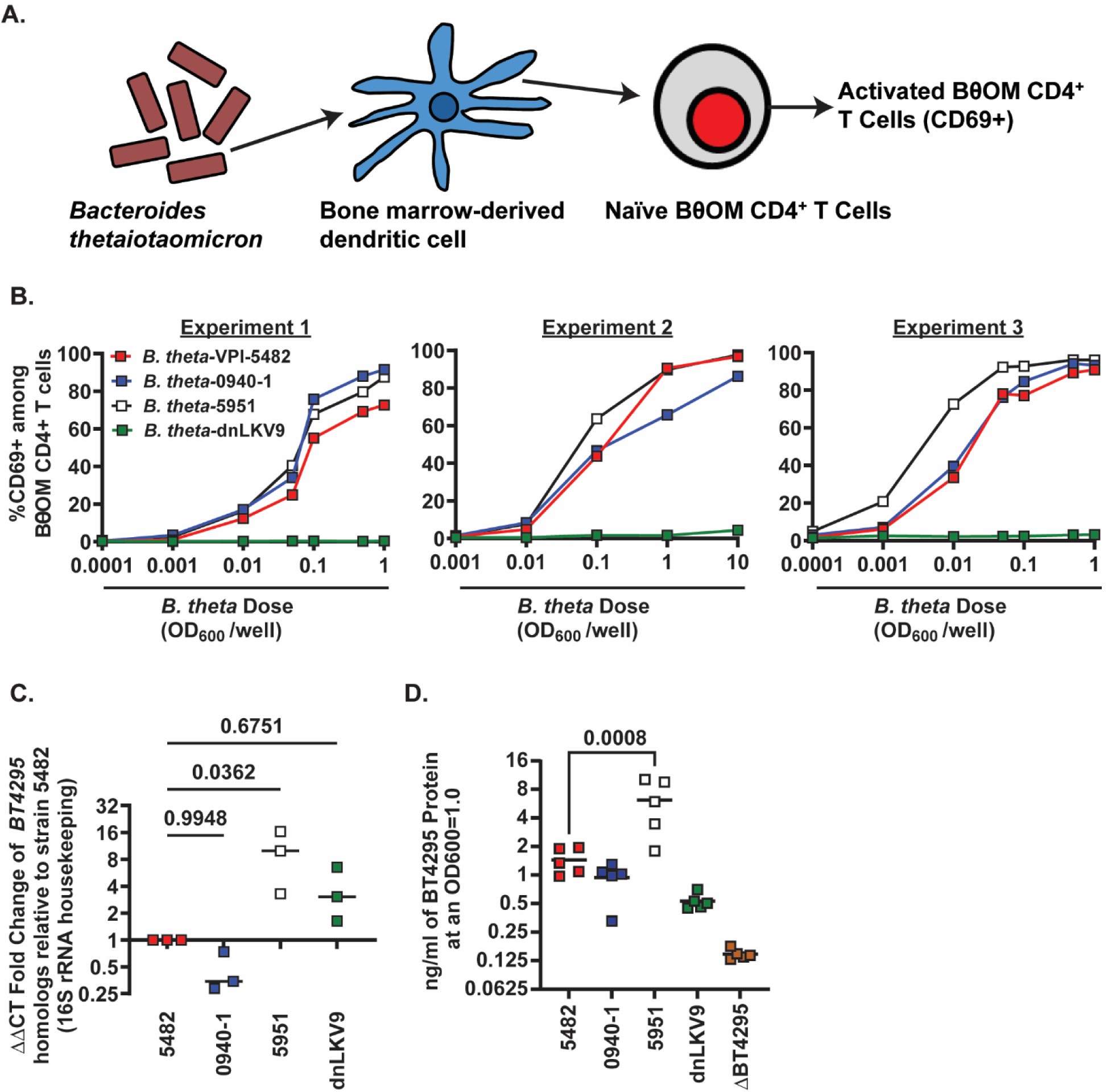
– Select *B. theta* strains evade recognition by *B. thetaiotaomicron*-specific BθOM CD4+ T cells. (A) Schematic of experimental scheme to assess activation of BθOM CD4+ T cells by strains of *B. theta*. (B) Bone-marrow derived dendritic cells were pre-loaded with the indicated doses of heat-killed strains of *B. thetaiotaomicron* for 24 hours and then mixed 1:1 with naïve BθOM CD4+ T cells. After 2 days CD69 expression was assessed by flow cytometry. Summary data of experiments described in (A) for the indicated strains of *B. theta.* Each data point represents a single technical replicate at the indicated dose, and three independent biological experiments are shown. (C) The expression of *BT4295* or its relevant homologs in the indicated strains of *B. theta* was assessed at stationary phase, using the 16S rRNA gene as a housekeeping gene to calculate relative expression. The graph shows the fold change in expression relative to expression in *B. theta*-VPI-5482 as calculated by the ΔΔCt method. Bars show the median and each point represents an individual biological replicate. Statistical significance was determined via one-way ANOVA with Dunnett’s multiple comparison test, comparisons made to *B. theta*-VPI-5482. (D) The expression of BT4295 protein in the indicated strains of *B. theta* was assessed at stationary phase using a custom anti-sera generated against recombinant BT4295. Bars show the median and each point represents an individual biological replicate. Statistical significance was determined via one-way ANOVA with Dunnett’s multiple comparison test, comparisons made to *B. theta*-VPI-5482. Background recognition of the anti-sera is shown by signal from *B. theta*^Δ^*^BT4295^*.

The expression of *BT4295* has been shown to be exquisitely sensitive to components of the media in which *B. theta* is grown [32]. To exclude the possibility that *B. theta*-dnLKV9 possessed the capacity to stimulate BθOM CD4+ T cells but simply had differential nutritional cues/requirements for induction of *BT4295* expression compared to the other strains tested, we measured the expression of *BT4925* by reverse transcriptase-quantitative PCR (RT-qPCR) in stationary phase (this time-point was chosen for measurement as this was the stage of the growth phase from which cells were harvested for use in stimulation assays). Although *B. theta*-dnLKV9 expressed lower *BT4295* transcript than *B. theta*-strain 5951 (a potent activator of BθOM CD4+ T cells), its expression was equivalent if not greater than that of *B. theta*-VPI-5482 (**Fig 1C**), thus excluding the possibility that a simple failure to transcribe the gene could be responsible for the failure to stimulate BθOM CD4+ T cells. In addition, we generated anti-BT4295 anti-sera and found that protein expression of BT4295 was lower relative to other strains (**Fig 1D**), suggesting that some feature of the protein in this strain was impaired.

To more comprehensively determine if *B. theta* dnLKV9 was impaired in activating BθOM CD4+ T cells rather than simply in the induction of the early activation marker CD69, we assessed expression of CD25, and secreted IL-2, both of which are induced upon T cell activation [57–59]. As with CD69 (**Fig 2A**), there were dose-dependent increases in CD25 expression (**Fig 2B**) and IL-2 secretion (**Fig 2C**) following stimulation with *B. theta* strains VPI-5482, 0940-1 and 5951, but not dnLKV9. Collectively, as these data show defective activation of both proximal (CD69) and distal (IL-2) markers of CD4+ T cell activation, they further establish that *B. theta* dnLKV9 is impaired in the activation of BθOM CD4+ T cells.

**Figure 2.**
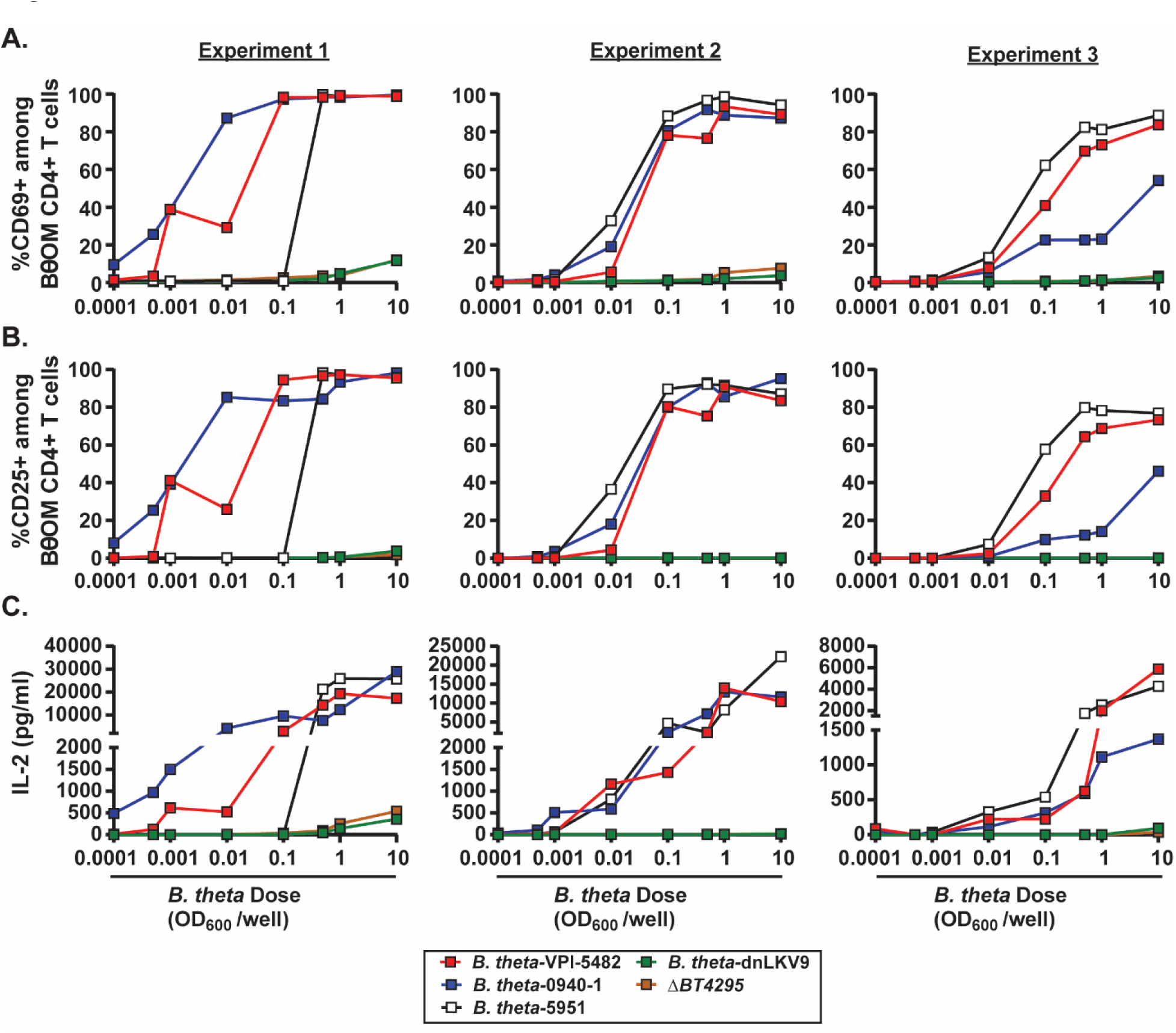
– Evasion of *B. thetaiotaomicron*-specific BθOM CD4+ T cells by *B. theta* strains impacts many stages of activation. Bone-marrow derived dendritic cells were pre-loaded with the indicated doses of heat-killed strains of *B. thetaiotaomicron* for 24 hours and then mixed 1:1 with naïve BθOM CD4+ T cells. (A-C) After 2 days the expression of cell surface (A) CD69 or (B) CD25 was assessed by flow cytometry, or (C) IL-2 secretion into culture supernatant was assessed by ELISA. Each data point represents a single technical replicate at the indicated dose, and three independent biological experiments are shown.

To further test whether *B. theta*-dnLKV9 truly lacked the capacity to stimulate BθOM CD4+ T cells, we utilized an *in vivo* model system whereby germ-free RAG1−/− mice were monocolonized with *B. theta*-VPI-5482 or *B. theta*-dnLKV9. After ∼2 weeks, all mice were adoptively co-transferred with a mixture of unfractionated wild-type CD4+ T cells from a CD45.1-CD45.2+ donor and naïve BθOM CD4+ T cells from a RAG1−/−CD45.1+CD45.2-donor (∼95% wild-type and ∼5% BθOM) (**Fig 3A)**. The co-transfer of wild-type cells: (i) allowed us to compare the fitness of BθOM CD4+ T cells relative to a reference population that should be unaffected by capacity of the different strains to stimulate BθOM cells, thus providing an internal control for each mouse, (ii) provided a control that allowed us to distinguish between issues affecting reconstitution of transferred cells from direct results of antigen availability (the wild-type cells should reconstitute mice regardless of the colonized microbes while the BθOM CD4+ T cells depend on the presence of *BT4295*), (iii) helped to limit issues associated with an overabundance of single clonotype [60–63], and (iv) limited the potential that BθOM CD4+ T cells could induce inflammatory responses against *B. theta* by providing a pool of regulatory T cells that would limit such responses. ∼3 weeks after cell transfer, BθOM (TCRVβ12+CD45.2-CD4+ T cells) and wild-type CD4+ T cells (CD45.2+ CD4+ T cells) were quantified in the spleen and colon of recipient mice. Strikingly, while wild-type cells readily accumulated in all mice in both spleen and colon, *B. theta*-VPI-5482 colonized mice had significantly greater accumulation of BθOM cells compared to *B. theta*-dnLKV9 (**Fig 3B-3E and Fig S2**). This was true both with respect to the proportion of BθOM cells within the CD4+ T cell pool (**Fig 3B-3D**) as well as the total number of BθOM CD4+ T cells (**Fig 3E)**, yet there were equivalent levels of colonization by the two strains (**Fig 3F**). Despite the impaired accumulation of BθOM CD4+ T cells, the total number of colonic CD4+ T cells (encompassing both wild-type and BθOM CD4+ T cells) did not differ between the mice colonized with the two strains (**Fig 3G**), suggesting that the effects observed with BθOM CD4+ T cells were not simply attributable to general differences in reconstitution dynamics based on the strains. Thus, these data reinforce the results obtained using our *in vitro* system and reveal that *B. theta*-dnLKV9 is significantly impaired in its capacity to stimulate BθOM CD4+ T cells relative to other strains of *B. theta*.

**Figure 3.**
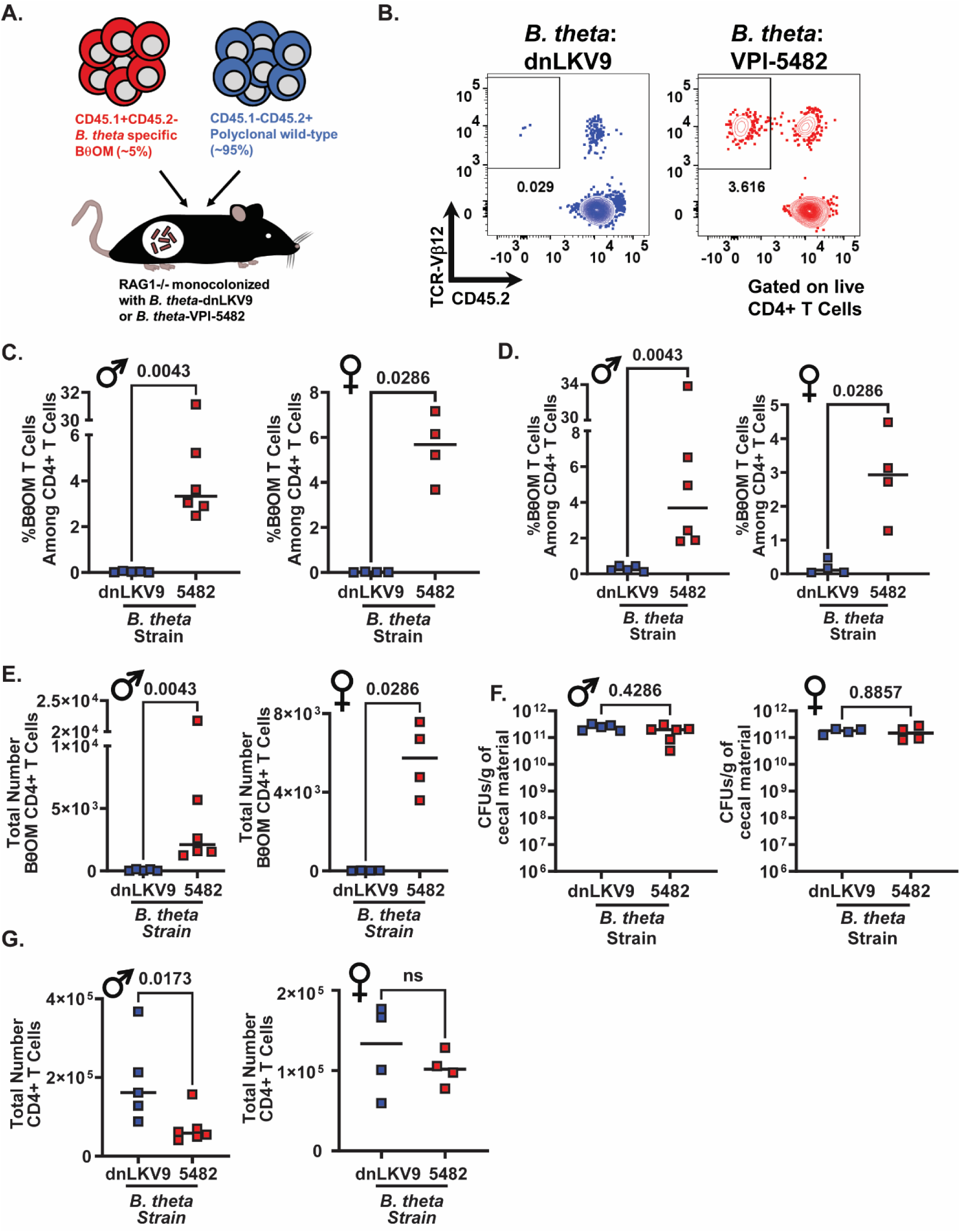
– B. *theta*-dnLKV9 evades recognition by BθOM CD4+ T cells *in vivo*. Naïve BθOM CD4+ T cells were mixed with unfractionated wild-type polyclonal CD4+ T cells and adoptively transferred to gnotobiotic RAG1−/− mice that had been colonized for ∼2 weeks with the indicated strain of *B. theta*. ∼3 weeks later the accumulation of BθOM CD4+ T cells was quantified (A) Schematic outlining experimental design for cell transfer. (B) Representative flow cytometry plots showing BθOM CD4+ T cell accumulation in the colon of the indicated groups of recipient mice (data shown is from male mice). (C) Graph shows the proportion of BθOM CD4+ T cells among all CD4+ T cells in the colon for both male and female recipients ∼3 weeks post cell transfer. (D) Graph shows the proportion of BθOM CD4+ T cells among all CD4+ T cells in the spleen for both male and female recipients ∼3 weeks post cell transfer. (E) Graph shows the total number of BθOM CD4+ T cells in the colon of male and female recipients ∼3 weeks post cell transfer. (F) Graph shows the colonization levels of *B. theta* as quantified by enumeration of colony forming units (CFU) in the cecum ∼5 weeks post colonization, ∼3 weeks post cell transfer. (G) Graph shows the total number of CD4+ T cells, encompassing BθOM and wild-type CD4+ T cells in the colon of both male and female recipient mice ∼3 weeks post cell transfer. Bars show the median and data points represent individual mice. Male and female cohorts represent two independent experiments. Statistical significance was determined via Mann-Whitney test. The data from one mouse was excluded from analysis due to technical issues during injection. The decision to exclude this data was taken at the time of injection prior to any knowledge of the outcome of the data. The exclusion of this data point did not affect conclusions drawn from the data. P values <0.05 were considered to be statistically significant. Symbols indicate the sex of the mice used: **♂** for male and **♀** for female.

### *B. theta* strain dnLKV9 harbors a variant copy of *BT4295*

We reasoned that the strain-level variation observed could reflect a strain-specific feature that modulates immunogenicity, or that there may be alterations to the antigen itself or the epitope within it that impacted expression of the protein or recognition by BθOM CD4+ T cells respectively. To explore this further, we examined the sequence of the *BT4295* homolog encoded by *B. theta*-dnLKV9 (locus tag, C799_02394). As with *B. theta*-VPI-5482, the canonical epitope recognized by the BθOM TCR (EEFNLPTTNGGHAT) was found in *B. theta*-dnLKV9 (**Fig 4**), excluding the possibility that a lack of the epitope could explain the failure to be recognized by BθOM CD4+ T cells. However, we noted a mutation downstream of the epitope that introduced a premature stop codon, and this mutation was not evident in any of the other strains analyzed (**Fig 4**). Were the protein to be translated and processed as normal, the epitope would be expressed and would be expected to stimulate BθOM CD4+ T cells. Thus, we rationalized that this premature stop codon impaired the proper maturation of the protein or its processing for presentation of the epitope on MHCII. We first sought to revert this mutation within *B. theta*-dnLKV9 such that the premature stop codon was removed, but our efforts to genetically modify *B. theta*-dnLKV9 proved unsuccessful. To overcome these challenges, we utilized a system whereby wild-type versions or variants of *BT4295* could be reconstituted in a *B. theta*^Δ*tdk*Δ*BT4295*^ background (on the VPI-5482 background; the Δ*tdk* background facilitates counterselection during construction of mutants of interest). Using the *in vitro* assay system described above, we found that *B. theta*^Δ*tdk*Δ*BT4295*^ reconstituted with wild-type *BT4295* (*B. theta*^Δ*tdk*Δ*BT4295*::*BT4295*^) drove CD69 expression in a dose-dependent manner in BθOM CD4+ T cells in our *in vitro* system relative to *B. theta*^Δ*tdk*Δ*BT4295*^ (**Fig 5A**), thus validating the approach. To determine if the premature stop codon found in *B. theta*-dnLKV9 could explain the failure to stimulate BθOM CD4+ T cells, we reconstituted *B. theta*^Δ*tdk*Δ*BT4295*^ with a truncated form of *BT4295* that mimicked the protein that would be formed due to the early stop codon in *B. theta*-dnLKV9 (*B. theta*^Δ*tdk*Δ*BT4295*::dnLKV9_trunc^) (note, although other amino acid differences exist between *BT4295* in *B. theta*-VPI-5482 and its homolog in *B. theta*-dnLKV9, this construct represented the *B. theta*-VPI-5482 form modified only to introduce the premature stop codon found in *B. theta*-dnLKV9; amino acids downstream of the stop codon were deleted). Strikingly, as with *B. theta*-dnLKV9, *B. theta*^Δ*tdk*Δ*BT4295*::dnLKV9_trunc^ failed to stimulate BθOM CD4+ T cell activation, as assessed by CD69 (**Fig 5B**) or CD25 (**Fig 5C**) expression, and secretion of IL-2 (**Fig 5D**). Interestingly, expression of BT4295 protein was reduced upon introduction of the premature stop codon (**Fig 5E**), mirroring the lower expression observed in *B. theta*-dnLKV9. Thus, although additional mechanisms may contribute to immune evasion in *B. theta*-dnLKV9, these data demonstrate that the introduction of a premature stop codon as found in *B. theta*-dnLKV9, in replacement of endogenous *BT4295* in *B. theta*-VPI-5482 is sufficient to confer the inability to activate BθOM CD4+ T cells, likely due to impaired expression. Moreover, these data reveal that both *B. theta*-dnLKV9 and *B. theta*^Δ*tdk*Δ*BT4295*::dnLKV9^ were as defective as *_B. theta_*^Δ*tdk*Δ*BT4295*^.

**Figure 4.**
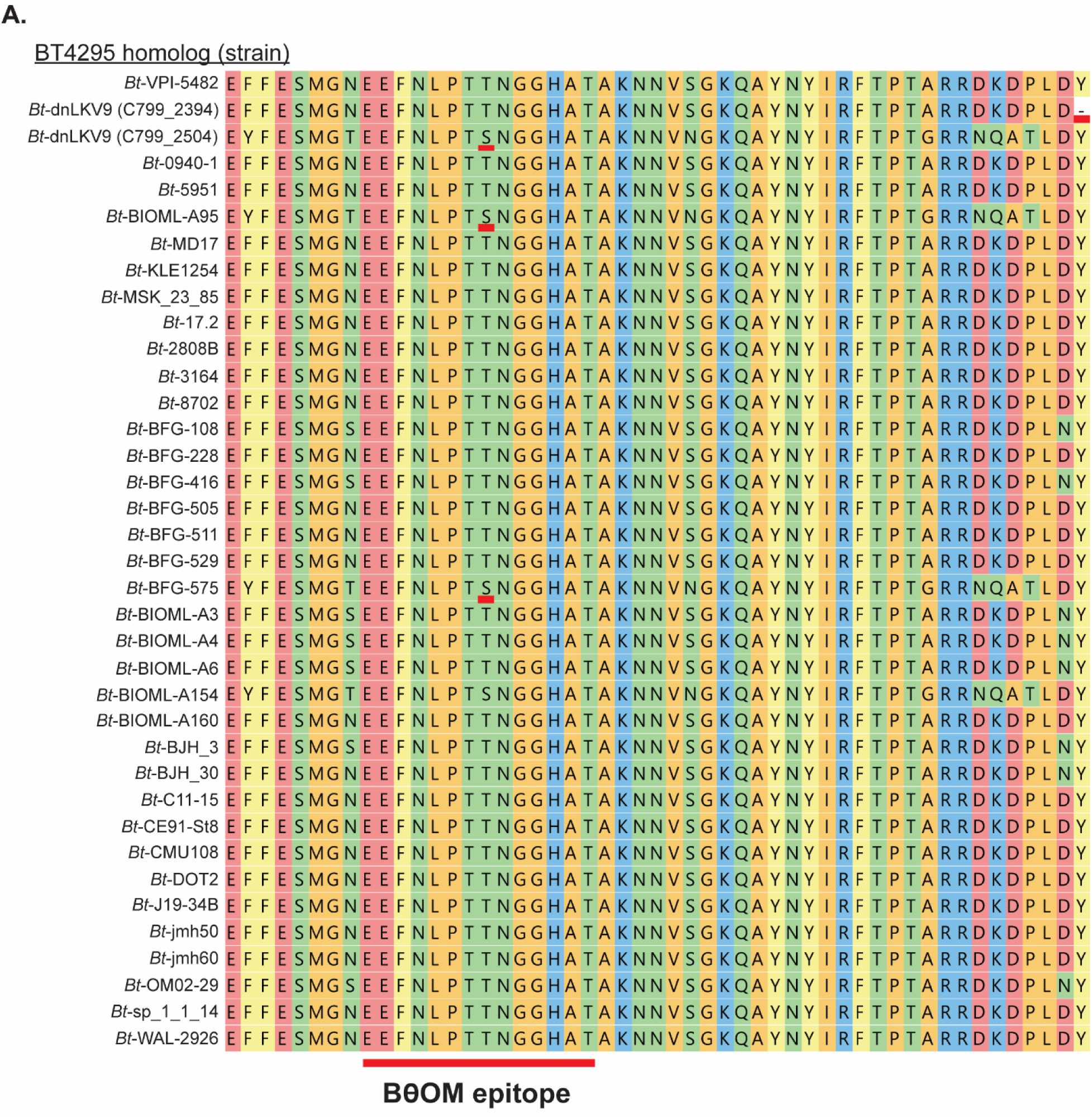
– Bioinformatic analyses reveal variation in BT4295 across strains of *B. theta*. The amino acid sequence of BT4925 and its homologs from the indicated strains of *B. theta* was extracted from publicly available genome sequences and aligned to facilitate comparisons between the different strains. The illustration shows the region of BT4295 that contains the canonical form of the epitope recognized by BθOM CD4+ T cells and the region immediately downstream. The presence of an alteration in epitope sequence (T548S) for *B. theta*-dnLKV9 (C799_2504), *B. theta*-BIOML-A95 and *B. theta*-BFG-575 is underlined in red. The existence of a premature stop codon downstream of the epitope in *B. theta*-dnLKV9 (C799_2394) (marked as a “-“), but not other strains, is also highlighted in red. Note, *B. theta*-dnLKV9 (C799_2504) and *B. theta*-dnLKV9 (C799_2394) represent proteins encoded by two different homologs of *BT4295* in *B. theta*-dnLKV9.

**Figure 5.**
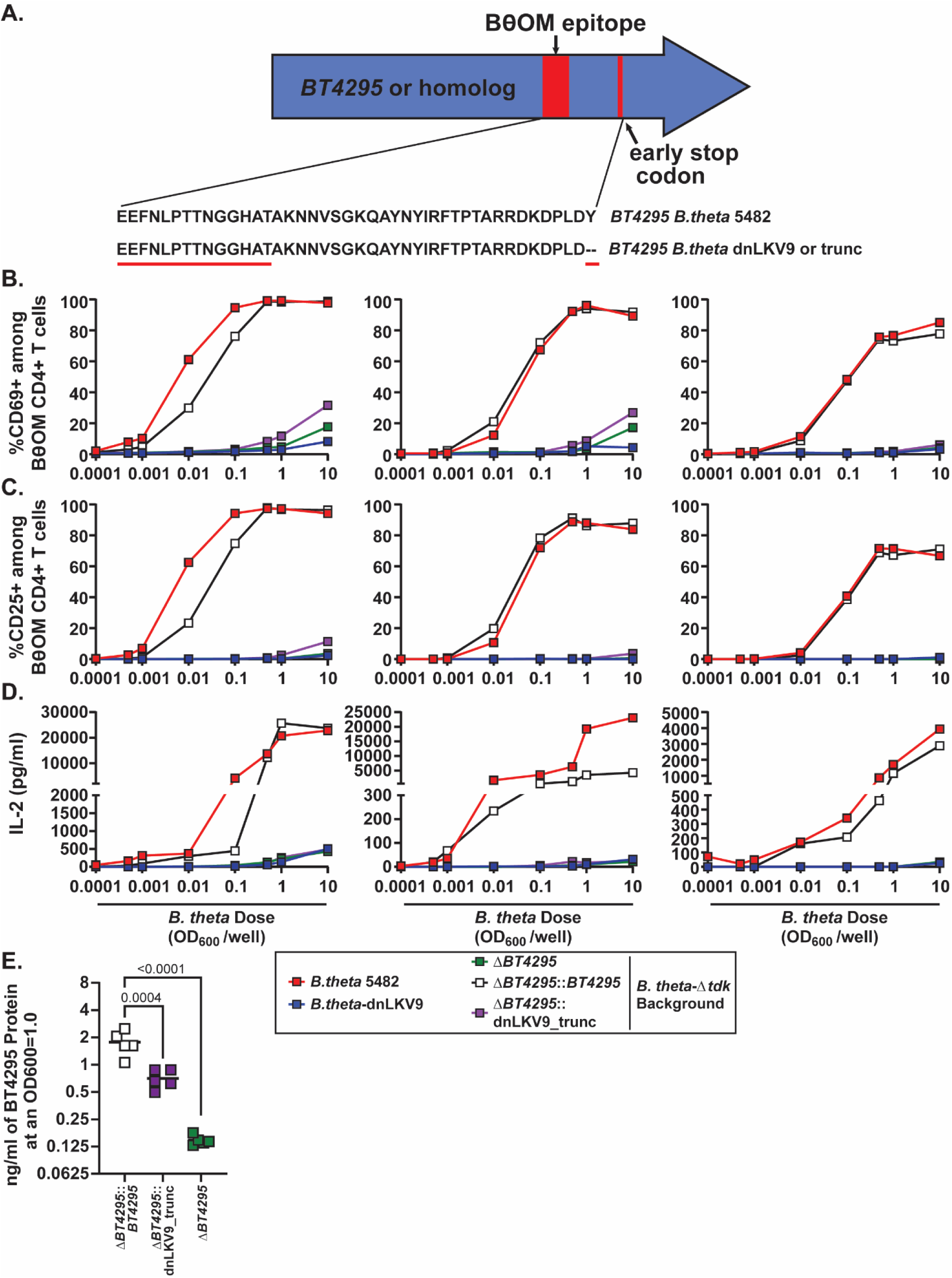
– A premature stop codon within BT4295 ablates recognition by BθOM CD4+ T cells. (A) Schematic of a premature stop codon that was introduced into *B. theta*-VPI-5482^Δ*tdk*Δ*BT4295*^ that mimicked that found in *B. theta*-dnLKV9 (*B. theta*-VPI-5482^Δ*tdk*Δ*BT4295*::dnLKV9_trunc^). Note, all genetically modified strains were generated on a *B. theta*-VPI-5482^Δ*tdk*Δ*BT4295*^ background. (B-D) Bone-marrow derived dendritic cells were pre-loaded with varying doses of heat-killed *B. theta*-VPI-5482^Δ*tdk*Δ*BT4295*::dnLKV9_trunc^ or the indicated controls for 24 hours following which they were mixed 1:1 with naïve BθOM CD4+ T cells. After 2 days cell surface (B) CD69 or (C) CD25 expression was assessed by flow cytometry. (D) IL-2 secreted into culture supernatant was assessed by ELISA. (E) BT4295 protein expression in the indicated strains of *B. theta* was assessed at stationary phase using a custom anti-sera generated against recombinant BT4295. This analysis was performed concurrently with the data in Figure 1D and thus the expression depicted for *B. theta*^Δ*tdk*Δ*BT4295*^ represents the same data presented in that figure. Each data point represents a single technical replicate at the indicated dose, and summary data for three independent biological experiments across a range of doses of *B. thetaiotaomicron* are shown (A-D), or bar shows the mean and each point represents individual biological replicates (E). Statistical significance was assessed by one-way ANOVA with Dunnett’s multiple comparison test.

### *B. theta*-dnLKV9 harbors an altered version of the epitope recognized by BθOM CD4+ T cells that facilitates immune evasion

During our efforts to identify the *BT4295* homolog in *B. theta*-dnLKV9 we noted the existence of a second gene (locus tag, C799_02504) coding for a product with significant homology to *BT4295* that was present in a distinct locus (**Fig 4**; depicted in the figure as “dnLKV9 (T548S)”). Interestingly, this homolog is not present in *B. theta*-VPI-5482. Notably, and in contrast with the homolog described above, this additional copy of the gene lacked a premature stop codon, and thus should have allowed for stimulation of BθOM CD4+ T cells. This prompted us to search for other mutations that could explain the failure of strain dnLKV9 to activate BθOM CD4+ T cells, leading to the finding that it contained an altered version of the canonical epitope with a T548S substitution when compared to the epitope found in *B. theta*-VPI-5482 (**Fig 4**).

To determine if this additional copy represented an altered peptide ligand [64] for BθOM CD4+ T cells we first tested the capacity of synthetic peptides representing the wild-type form of the epitope (EEFNLPTTNGGHAT) or the T548S substituted form (EEFNLPT<u>S</u>NGGHAT; substitution position underlined) to stimulate upregulation of CD69 *in vitro*. Strikingly, the T548S peptide was severely impaired in its capacity to drive enhanced activation of BθOM CD4+ T cells (as assessed by expression of CD69 and CD25, and secretion of IL-2) using the *in vitro* assay system described above. However, at the highest doses tested it was able to promote a maximal CD69 (**Fig 6A**) and CD25 (**Fig 6B**) expression, as well as IL-2 secretion (**Fig 6C**) at or nearly equivalent to the maximum induced by the wild-type form of the epitope (note, at the highest doses of WT peptide significant down-regulation of the TCR was observed and thus some cells were not captured in our gating strategy; inclusion of TCRβ-events showed almost identical results, not shown). Interestingly, the T548S form did not ablate the capacity of the wild-type form of the epitope to drive BθOM CD4+ T cell activation when BMDC were loaded with equivalent concentrations of both wild-type and T548S forms of the peptide (although minor effects were observed in one experiment) (**Fig 6A-C**). Thus, these data identify the existence of an altered peptide ligand (APL) of the canonical epitope, and although of lower potency, the T548S substituted form of the epitope did not possess strong BθOM inhibitory capacity, suggesting that it lacks the antagonist function reported for many APL [65].

**Figure 6.**
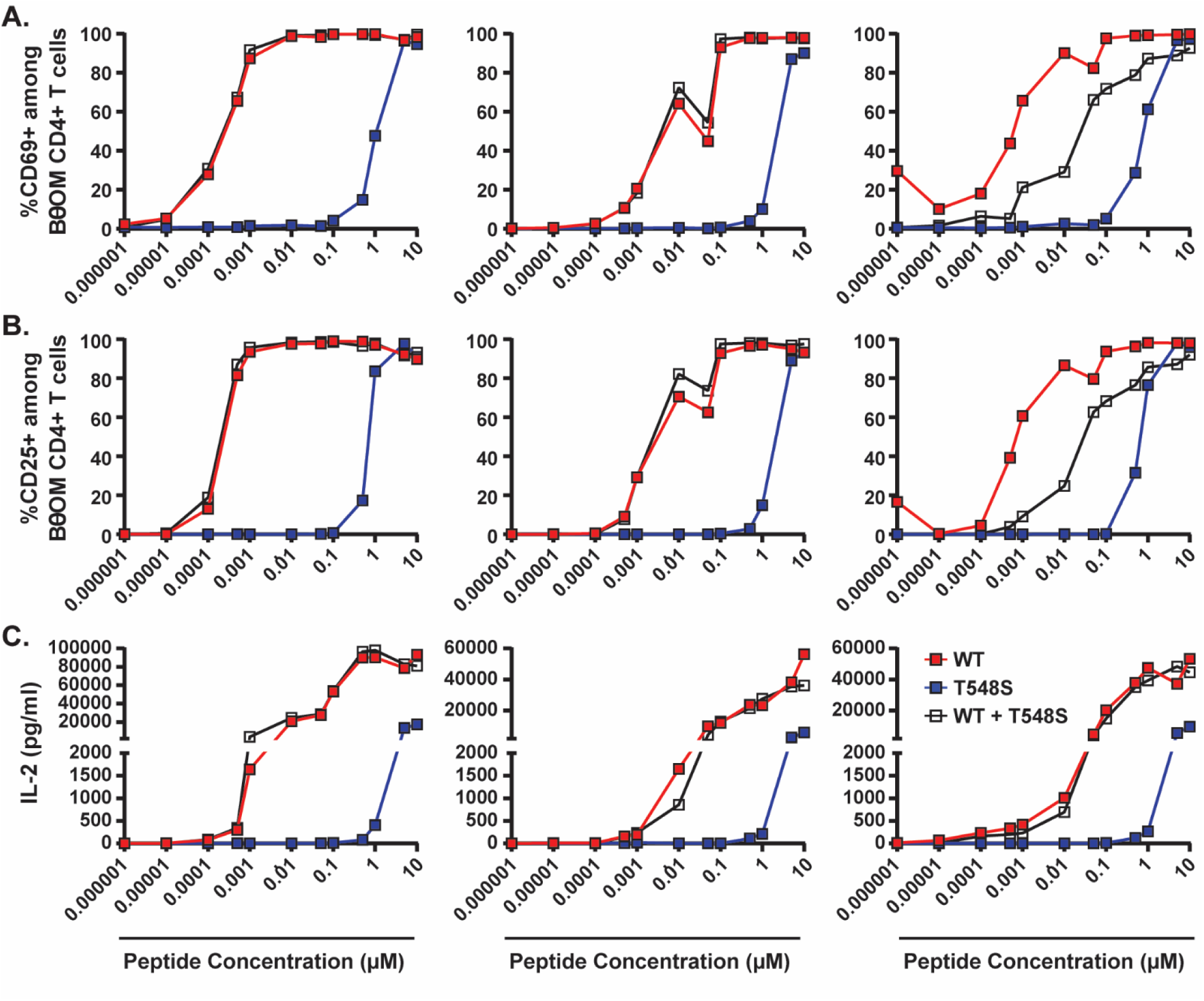
– B. *theta*-dnLKV9 harbors an altered peptide ligand for BθOM CD4^+^ T cells with reduced potency. Bone-marrow derived dendritic cells were loaded with varying doses of synthetic peptides representing the canonical form of the epitope for BθOM CD4+ T cells found in *B. theta*-VPI-5482, a mutated form of the epitope found in *B. theta*-dnLKV9 (T548S), or a mixture of both peptides, for 24 hours. Following this, the cells were mixed 1:1 with naïve BθOM CD4+ T cells and were incubated for 2 days. Cell surface (A) CD69 or (B) CD25 expression was then assessed by flow cytometry, (C) IL-2 secreted into culture supernatant was assessed by ELISA. Each point represents a single technical replicate at the indicated dose. The data shown are show data from three independent experiments that assessed BθOM CD4+ T cell activation across a range of doses of peptides (A). Data for CD69 are representative of a total of six experiments.

Next, we again leveraged our capacity to introduce modified forms of *BT4295* to *BT4295*-deficient *B. theta*-VPI-5482 and generated a form that expressed *BT4295* with a T548S substitution, *B. theta*^Δ*tdk*Δ*BT4295*::T548S^. In addition, to determine if position 548 was a critical residue for interaction of the BθOM TCR with *BT4295*, we generated additional mutants at this position, specifically, T548A and T548V, and assessed their impact on activation of BθOM CD4+ T cells in our *in vitro* assay system through assessment of CD69 and CD25, and secretion of IL-2. Similarly to the synthetic peptide, we found that *B. theta*^Δ*tdk*Δ*BT4295*::T548S^ displayed a severely impaired capacity to stimulate BθOM CD4+ T cells relative to a *B. theta*^Δ*tdk*Δ*BT4295*::*BT4295*^ control, but could efficiently activate these cells at higher doses than required for *B. theta*^Δ*tdk*Δ*BT4295*::*BT4295*^ (**7A-C**) in keeping with the idea that this mutation alters its potency as an agonist rather than ablating recognition completely. The T548A substitution also severely impaired activation of BθOM CD4+ T cells as recently established by others [66], but the T548V mutation had little effect or even enhanced activation (**Fig 7A-C**). The T548S form of BT4295 was expressed at equivalent or greater levels than the wild-type form, thus pointing to impaired recognition of this form of the epitope by BθOM CD4+ T cells (**Fig 7D**). Given this, the fact that the synthetic peptide produced similar results, and the magnitude of reduction in activating capacity, this expression difference is not likely the reason for impaired potency. Thus, our data shows that position 548 appears to be a critical residue in BT4295 for recognition by BθOM CD4+ T cells, and modulation at this position has varying impacts on stimulation of BθOM CD4+ T cells depending on the nature of the modification.

**Figure 7.**
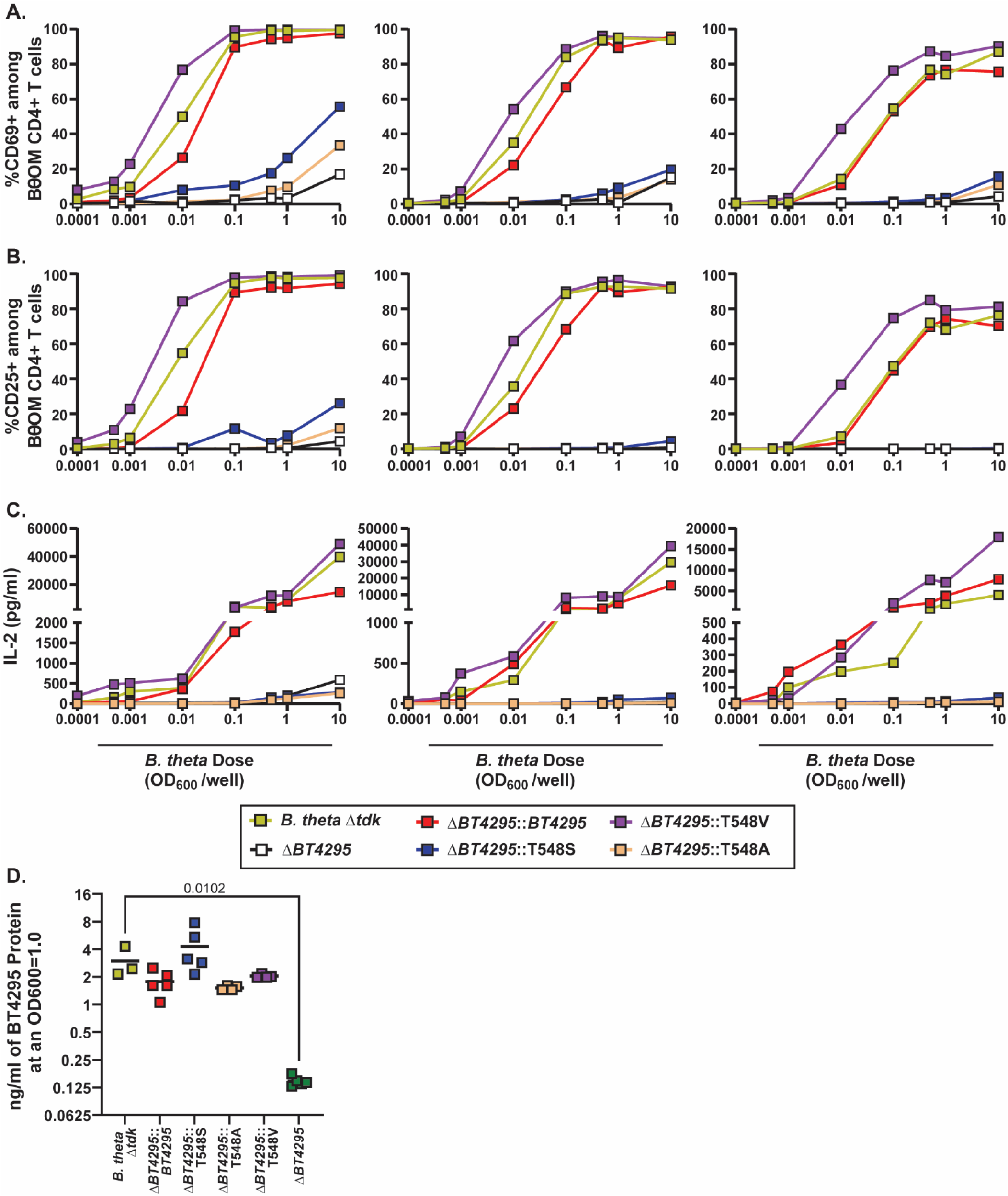
– B. *theta*-dnLKV9 harbors an altered peptide ligand for BθOM CD4^+^ T cells with reduced potency. Bone-marrow derived dendritic cells were loaded with varying doses of heat-killed *B. theta*-VPI-5482^Δ*tdk*Δ*BT4295*^ that had been modified to express the indicated variants of the epitope recognized by BθOM CD4+ T cells, for 24 hours. Following this, the cells were mixed 1:1 with naïve BθOM CD4+ T cells and were incubated for 2 days. (A-C) Cell surface (A) CD69 or (B) CD25 expression was then assessed by flow cytometry, or (C) IL-2 secreted into culture supernatant was assessed by ELISA. (D) BT4295 protein expression was assessed by ELISA using a custom rabbit anti-sera. This analysis was performed concurrently with the data in Figure 1D and Figure 5E, and thus the expression depicted for *B. theta*^Δ*tdk*Δ*BT4295*^ and *B. theta*^Δ*tdk*Δ*BT4295::BT4295*^ represents the same data presented in those figures. Each point represents a single technical replicate at the indicated dose, (A-C). The data shown represent three independent experiments that assessed BθOM CD4+ T cell activation across a range of doses of peptides. Data for CD69 are representative of six independent experiments. In (D) the bar represents the mean and each data point represents an individual biological replicate. Statistical significance was assessed by one-way ANOVA with Dunnett’s multiple comparison test.

Modulation of the agonist potency has been linked to alterations to both the magnitude and quality of the T cell response that ensues following T cell stimulation [67–75]. Given that the T548S mutation impaired but did not abolish the capacity to stimulate BθOM CD4+ T cells we wanted to determine if the mutation impacted the accumulation of BθOM CD4+ T cells and/or their phenotype *in vivo*. We therefore used an experimental setup as described above, where germ-free RAG1−/− mice were monocolonized with *B. theta*^Δ*tdk*Δ*BT4295*::*BT4295*^ or *B. theta*^Δ*tdk*Δ*BT4295*::T548S^ for ∼2 weeks, following which all mice were adoptively transferred with a mix of naïve BθOM CD45.1+CD45.2-CD4+ T cells and bulk wild-type CD45.1-CD45.2+ CD4+ T cells (**Fig 8A**). ∼3 weeks post transfer, the accumulation of BθOM CD4+ T cells was assessed in the spleen and colon (**Fig S2B**). BθOM CD4+ T cells were massively impaired in their capacity to expand in mice colonized with *B. theta*^Δ*tdk*Δ*BT4295*::T548S^, both in terms of proportion of the colonic (**Fig 8B and 8C**) and splenic (**Fig 8D**) CD4+ T cell pool as well as the total number of BθOM cells that accumulated in the colon (**Fig 8E**), relative to mice colonized with the control *B. theta*^Δ*tdk*Δ*BT4295*::*BT4295*^ that expressed the wild-type form of the epitope. Indeed, the numbers of BθOM CD4+ T cells were so low that accurate phenotyping of their differentiation profile was not possible. No differences were observed in colonization capacity between the two strains (**Fig 8F**) suggesting that impaired colonization does not contribute to this phenotype. The total number of CD4+ T cells that accumulated in the colon was also not significantly different between the two groups (**Fig 8G**), suggesting that the impairment in BθOM CD4+ T cells was a specific effect and not simply a more generalized feature that also affected BθOM CD4+ T cell reconstitution. Thus, despite the fact that it can stimulate BθOM CD4+ T cells *in vitro*, the T548S form of the epitope that is expressed in *B. theta*-dnLKV9 is also not sufficient to restore its capacity to stimulate the accumulation of BθOM CD4+ T cells in the intestine, likely because it does not provide adequate signal through the TCR at the doses available *in vivo*.

**Figure 8.**
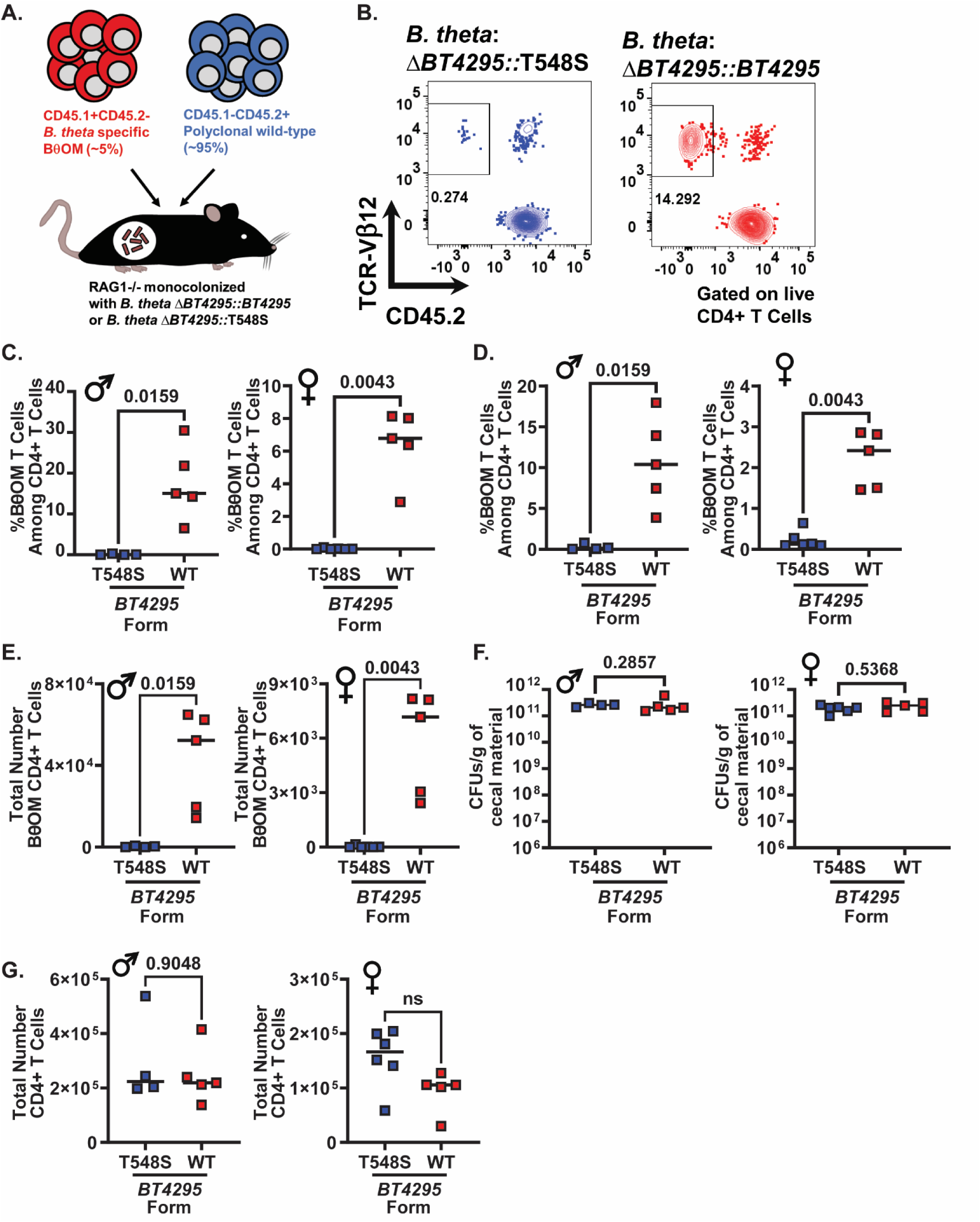
– An altered peptide ligand for BθOM CD4^+^ T cells in *B. theta*-dnLKV9 mediates immune evasion *in vivo*. Naïve BθOM CD4+ T cells were mixed with unfractionated wild-type polyclonal CD4+ T cells and adoptively transferred to gnotobiotic RAG1−/− mice that had been colonized for ∼2 weeks with the indicated strain of *B. theta*. ∼3 weeks later the accumulation of BθOM CD4+ T cells were quantified (A) Schematic outlining experimental design for cell transfer. (B) Representative flow cytometry plots showing BθOM CD4+ T cell accumulation in the colon of the indicated groups of recipient mice (data shown is from male mice). (C) Graph shows the proportion of BθOM CD4+ T cells among all CD4+ T cells in the colon for both male and female recipients ∼3 weeks post cell transfer. (D) Graph shows the proportion of BθOM CD4+ T cells among all CD4+ T cells in the spleen for both male and female recipients ∼3 weeks post cell transfer. (E) Graph shows the total number of BθOM CD4+ T cells in the colon of male and female recipients ∼3 weeks post cell transfer. (F) Graph shows the colonization levels of *B. theta* as quantified by enumeration of colony forming units (CFU) in the cecum ∼5 weeks post colonization, ∼3 weeks post cell transfer. (G) Graph shows the total number of CD4+ T cells, encompassing BθOM and wild-type CD4+ T cells in the colon of both male and female recipient mice ∼3 weeks post cell transfer. Bars show the median and data points represent individual mice. Male and female cohorts represent two independent experiments. Statistical significance was determined via Mann-Whitney test. P values <0.05 were considered to be statistically significant. Symbols indicate the sex of the mice used: **♂** for male and **♀** for female.

## Discussion

Here we investigated whether a species-specific CD4+ T cell clone, as represented by CD4+ T cells from BθOM TCR transgenic mice, could uniformly respond to multiple members of the *Bacteroides thetaiotaomicron* species or if particular strains could evade recognition. We found that while many tested strains were recognized by BθOM CD4+ T cells leading to their activation equivalent to the type strain *B. theta*-VPI-5482, one strain, *B. theta*-dnLKV9, was severely impaired in its capacity to activate these cells, both *in vitro* and *in vivo* in gnotobiotic mice. Importantly, this evasion of recognition could not simply be attributed to a lack of the gene encoding the activating epitope, *BT4295*. We identified that *B. theta*-dnLKV9 harbored a premature stop codon within its homolog of *BT4295* that was downstream of the epitope, and by reconstructing this mutation in *B. theta*-VPI-5482 that lacked its own copy of *BT4295*, showed that this mutation was sufficient to ablate recognition, thus conferring evasion of BθOM CD4+ T cells to a permissive strain. We further identified an additional homolog of *BT4295* that was present in *B. theta*-dnLKV9 but not *B. theta*-VPI-5482. This copy harbored an altered version of the epitope, with a T548S amino acid substitution that led to a significant reduction in the capacity to stimulate BθOM CD4+ T cells both *in vitro* and *in vivo* in gnotobiotic mice. Thus, our data uncover strain-specific modifications to an immunodominant CD4+ T cell antigen in *B. theta* that facilitate evasion of recognition by select CD4+ T cells. Our data point to mechanisms through which individual strains can evade recognition of T cell clones directed against antigens they express and point to a mechanism through which individualized pools of CD4+ T cells for strains can develop. Such a tailored response could explain how intestinal CD4+ T cells avoid inappropriate and damaging responses directed towards symbionts that have been generated in response to pathogenic isolates with the same epitope.

Several TCR transgenic mice have now been developed that are specific for microbiome-encoded epitopes [19, 26, 27, 30, 32, 76, 77]. These tools have provided pivotal insights into modulation of CD4+ T cell function by the microbiome, revealing that microbiome-specific CD4+ T cells can adopt a variety of cell fates following recognition [19, 20, 26, 27, 30, 32, 76], that vary depending both on the specific microbe that expresses the antigen [27], the presence or absence of inflammation [26, 30, 32, 78], and the nature of the other community members present [19, 48]. Despite these advances, it has remained largely unclear whether the observed responses are representative of the target species as a whole, or if strain-level variation exists, a feature of many interactions between hosts and pathogens that has only begun to be explored for microbiome-driven immune phenotypes [41–43, 45, 79–85]. Our studies suggest that while a given T cell clone may be responsive to many strains of a species, it cannot be assumed that a TCR of known specificity will respond to a microbe where this has not been empirically tested, even when the strains express the target epitope of interest. Together with an emerging body of work, our data suggest that strain-level variation is an important feature with respect to immune modulation by the microbiome that warrants more detailed study. Although it is important to highlight that the use of RAG-deficient and gnotobiotic mice can lead to greater activation of adoptively transferred microbiome-specific T cells than would be observed in more “natural” environments [48, 86], this provided the greatest opportunity for *B. theta*-dnLKV9 to activate BθOM CD4+ T cells, and thus any defects observed here are likely to be magnified in the context of lymphocyte replete hosts and/or mice born with complex microbiomes.

Prior studies have revealed the capacity of CD4+ T cell clones to respond to phylogenetically related microbes that share the target epitope, or even to modified forms of the epitope, providing a mechanism through which a broad array of microbes could be recognized by a single T cell clone [28, 29, 40]. More recently, it has been shown that the most abundant TCRs in the intestinal CD4+ T cell compartment react to microbiome-derived epitopes that are widely shared and highly expressed [40]. This model provides an explanation for how a limited number of CD4+ T cell clones can react to a broad array of microbes through focusing of the TCR repertoire on broadly and highly expressed antigens, supporting a notion of community-level epitope immunodominance. However, the model does not account for the need to tailor the immune response at the strain-level, due to the capacity of closely related strains to have opposing impact on host health [41–47]. Studies tracking the fate of an antigen from SFB have shown that when expressed in SFB, TCR transgenic cells specific for the target SFB-derived antigen adopt a Th17 fate in the intestine, but when the same antigen is expressed in *Listeria monocytogenes*, the cells adopt a Th1 fate [27]. Although this involved forced expression of an epitope that was not truly shared between SFB and *L. monocytogenes*, it demonstrated that distinct fates can be primed against the same epitope in the intestine, underscoring the need for responses to be tailored to the nature of the microbe from which they are derived rather than a uniform response irrespective of the impact of the source of the epitope on the host. Our data suggest that even if deleterious responses are primed against a shared epitope, individual strains that express the epitope may evolve strategies to evade recognition, or indeed simply acquire random mutations, that limit the activity of particular T cell clones against them. Collectively, these data add to the current model through which the regulation of intestinal CD4+ T cell responses is conceptualized, providing insights into how a CD4+ T cell compartment of a limited size balances the need for broad reactivity with the fine-level specificity needed to tailor responses in a strain-specific manner. Although much work remains to be done, we posit that for each individual strain the quality of the response will be dictated by the collective function of all the CD4+ T cells that recognize the strain, thus allowing a highly individualized output even where some of the clones within the pool have been polarized by a different strain. Under such a scenario, we envision that a collection of “public” T cell clones exist that are specific for a large number of community members that share an epitope in addition to “private” T cell clones that are highly strain-specific. Thus, for a commensal microbiome member with a closely related pathogen/pathobiont, T cell clones with inflammatory potential that are elicited by the pathogenic agent may also react to the commensal, but ultimately, the existence of sufficient anti-commensal T cell clones with anti-inflammatory potential will invoke tolerance to the commensal while allowing the function of protective inflammatory functions where necessary. Although it is clear that many other factors will contribute to the maintenance of immune tolerance, we hypothesize that this forms one of the layers of mutualism that limits deleterious anti-microbiome T cell responses in the intestine.

Although our data only provides evidence of evasion of a single TCR clone by one strain of bacteria, additional evidence for this phenomenon has been provided in a prior study of a distinct *B. theta*-reactive TCR [28]. Using a hybridoma-based approach, an epitope within BT0900 of *B. theta*-VPI-5482 was found to be shared with other *Bacteroides* species, specifically *B. finegoldii*, *B. ovatus*, and *B. caccae*, with a 100% match in epitope sequence across the different species. However, although both *B. finegoldii* and *B. ovatus* could activate this hybridoma, *B. caccae* lacked any capacity to do so above that of a *BT0900*-deletion mutant. While the mechanisms involved are undefined, these data reveal that evasion of individual CD4+ T cell clones, even when carrying an exact epitope match, is not simply a feature of *B. theta*-dnLKV9 and may represent a common feature among gut microbes.

Finally, our study identified the existence of a naturally occurring altered peptide ligand (APL) in an immunodominant epitope, represented by a T548S substitution in the epitope recognized by the *B. theta*-specific BθOM TCR. APLs have long been recognized as a means through which the strength of TCR stimulation and the quality of the ensuing response can be modulated [64, 65, 87–92]. As such, they have provided a means through which pathogens can disarm host-protective T cell responses, by antagonizing the TCR or altering the quality of the response [87–90]. As only a handful of microbiome-derived CD4+ T cell epitopes have been defined [26–29], and often few isolates of a species have complete genome sequences available, the prevalence of APLs in the gut microbiome has not been studied in detail, although the existence of altered epitopes that impaired recognition has been previously identified [29]. The APL identified here (EEFNLPT<u>S</u>NGGHAT) has weak stimulatory power *in vitro* and did not appear to prevent the stimulation when mixed with the native form of the epitope (EEFNLPTTNGGHAT), suggesting that it lacks the inhibitory capacity of many APLs [65, 93]. As naturally occurring APL of a flagellin-derived peptide from *Salmonella* stimulate lower production of interferon-γ from flagellin-specific SM1 cells relative to the native epitope [92] we were interested to determine if the T548S APL would skew the balance of different effector fates. In the intestine, BθOM CD4+ T cells adopt a FoxP3+ Treg fate as well as pro-inflammatory cytokine secreting potential that is limited by the developing Tregs [32]. However, as the T548S APL did not lead to appreciable accumulation of BθOM CD4+ T cells *in vivo*, we were unable to determine if this modulation shaped phenotypic effector outcomes.

Although our data do not provide a means through which the prevalence of such APL can be estimated, they suggest that such alterations may play an important role in shaping CD4+ T cell responses against individual strains in the gut. While it is tempting to speculate that the APL represents a means of escaping CD4+ T cell recognition in general, this cannot be reliably ascertained from our data. The BθOM TCR was isolated from popliteal lymph node CD4+ T cells from mice that had been immunized with *B. theta*-VPI-5482, and it is possible that the T548S version is equally immunogenic and would be the target of a strong CD4+ T cell response in the context of colonization/immunization with *B. theta*-dnLKV9. Indeed, *B. theta*-VPI-5482 may escape recognition by such cells directed against *B. theta*-dnLKV9. However, irrespective of whether the T548S APL represents an immune evasion strategy, or whether it simply represents an equally immunogenic variant recognized by a distinct T cell clone, our data show that *B. theta*-dnLKV9 can evade recognition by a CD4+ T cell clone that recognizes a wide-array of other *B. theta* strains. Thus, while it is likely that numerous CD4+ T cell clones that emerge following colonization with *B. theta*-dnLKV9 or *B. theta*-VPI-5482 would be strongly reactive to both strains, this is not a uniform process. Indeed, prior work has shown that the expression of *BT4295*, which encodes the epitope recognized by BθOM CD4+ T cells, is sensitive to dietary components [32], reinforcing the existence of myriad mechanisms through which recognition can be evaded. These data also highlight T548 as an important residue for BθOM CD4+ T cell recognition of the epitope, adding to previous findings that showed that T547 is a critical residue within the epitope [32]. Our work now extends our knowledge of the critical determinants within the epitope, and further reveals how it may represent a site whose modification can enhance or ablate recognition depending on the specific substitution.

We identified unexpected mutations in our BθOM mouse colony during the course of these studies, in keeping with prior work that has highlighted the potential for significant genetic differences in genetically modified lines of mice [94, 95]. While these mutations represent a limitation of our studies, and should be taken into account when interpreting our findings, all comparisons described assessed reactivity of BθOM CD4+ T cells to distinct strains/BT4295 antigen forms, and thus were equally affected by these mutations. Moreover, with respect to the loss of IL-10 specifically, given its immunosuppressive nature, it would be expected that cells would be easier to activate in the absence of a negative feedback loop from IL-10, yet we still observed significant impairment in activation in response to distinct forms of BT4295.

Collectively, our data show that CD4+ T cells should be considered as strain-specific and suggest the existence of public and private responses that balance to provide broad reactivity that maintains the strain-level discrimination needed to ensure the appropriate response is mounted against the numerous microbial strains resident in the gut microbiome. This adds to a growing body of work defining how immunity to *Bacteroides thetaiotaomicron* is mediated in steady state and disease [15, 18, 31, 32, 50, 56, 96], and underscores the need to study strain-level variation in gut microorganisms.

## Acknowledgments

We are indebted to the members of the Gnotobiotic Core facility at the Lerner Research Institute, especially Brandon Bakos, for help with execution of gnotobiotic experiments. We are extremely grateful to Prof. Thaddeus Stappenbeck for kindly providing us the *B. thetaiotaomicron*-dnLKV9 strain, Prof. Eric Martens for the provision of *BT4295* deficient *B. thetaiotaomicron*, Prof. Paul Allen for the provision of BθOM mice, Stephen Horvath and Darren Kreamalmeyer for advice regarding genotyping and breeding of BθOM mice, and Prof. Andrew Kau, Prof. Meng Wu and Dr. John Dekker for insightful comments on these studies.

## Author Contributions

R.W.P.G. designed and executed all experiments relating to the evasion of BθOM CD4+ T cells by various strains of *B. thetaiotaomicron*, generated the described genetically modified forms of *B. thetaiotaomicron*, analyzed and interpreted data, and helped write the manuscript. J.M.T., O.D.B., and M.J.E. assisted with the described experiments, and provided input on data interpretation. V.M. performed bioinformatic analyses of whole-genome sequencing of the BθOM mouse strain. P.P.A. helped to design the described experiments, interpret and analyze data, and helped to write the manuscript.

## Financial Support

R.W.P.G. is the recipient of NIH Loan Repayment Awards: LRP0000016021 and LRP0000045724. This work was supported by funds from (i) the Cleveland Clinic Foundation and R01DK126772 from the National Institute of Diabetes and Digestive and Kidney Diseases awarded to the lab of P.P.A, and (ii) R50CA293821 the National Cancer Institute to V.M.

## Conflicts

The authors do not have any financial conflicts of interest to declare.

## Data Availability

All data is available upon request from the corresponding author, Philip P. Ahern.

## Notes

### Competing Interest Statement

The authors have declared no competing interest.

### Summary of Updates

Phenotyping of the activation of CD4+ T cells has been extended to include assessment of CD25 and IL-2 expression, in addition to data presented regarding CD69 expression in the original submission. Furthermore, various forms of BT4295 expression are now assessed at the protein level.

