## Supplemental Figures for "Strain-level antigen variation facilitates immune evasion in *Bacteroides thetaiotaomicron*"

**Supplementary Figure 1-Gating strategies for purification of cells by flow cytometric based cell sorting**

Naïve BθOM CD4+ T cells for *in vivo* adoptive transfer experiments were purified by flow cytometric based cell sorting. Representative pre- and post- sort frequencies are shown from two independent experiments.

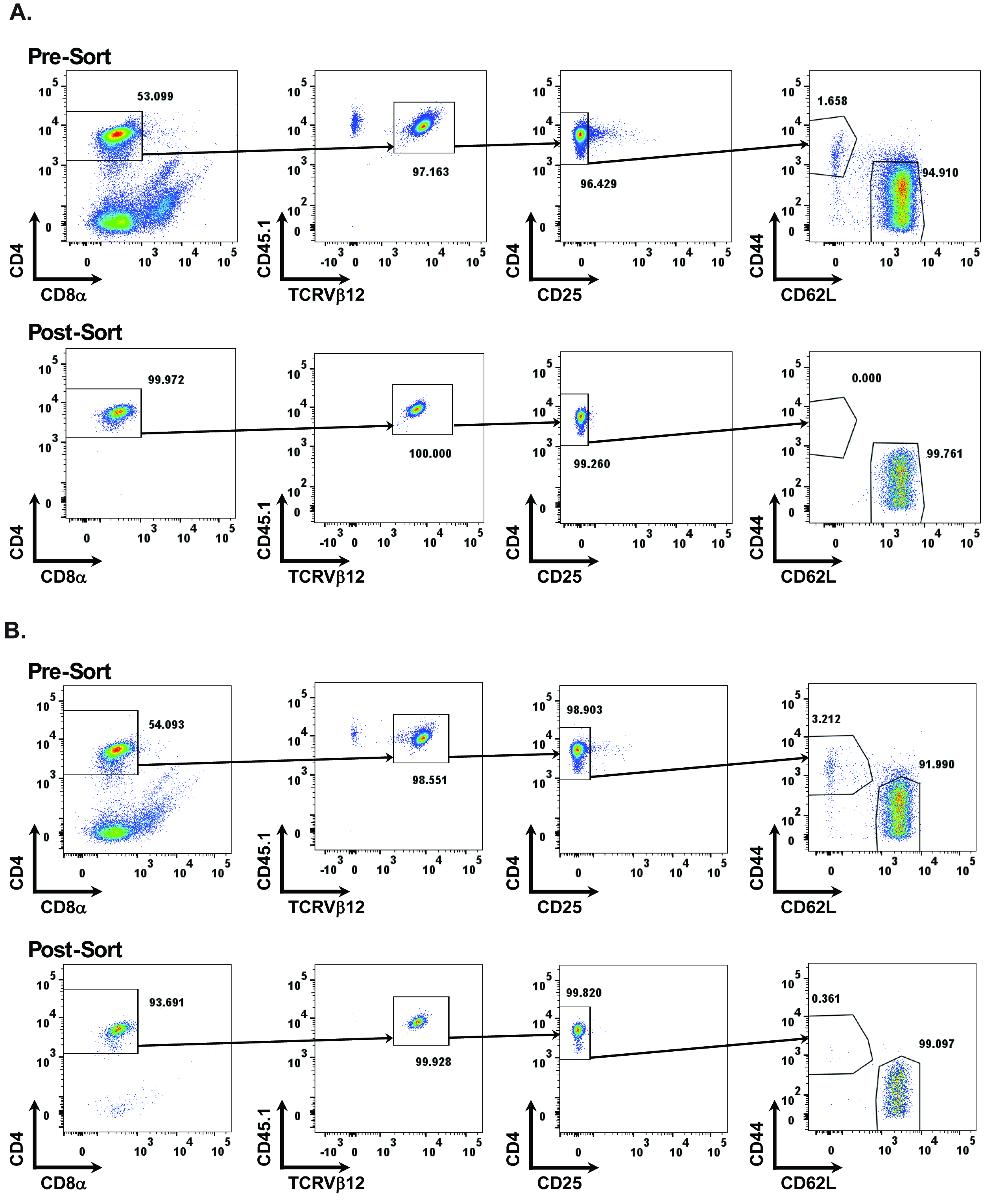

**Supplementary Figure 2-Gating strategies for analysis of BθOM CD4+ T cells by flow cytometry.**

BθOM CD4+ T cell activation following *in vitro* stimulation or following *in vivo* adoptive transfer was assessed via flow cytometry. Shown are representative gating strategies used to phenotype BθOM CD4+ T cells from these assays.

(A) Strategy for assessment of *in vitro* stimulation of BθOM CD4+ T cells
(B) Strategy for assessment of BθOM CD4+ T cells in experiments comparing *B. theta*-VPI-5482 and *B. theta*-dnLKV9 *in vivo*.
(C) Strategy for assessment of BθOM CD4+ T cells in experiments comparing *B. theta*^Δ^*^tdk^*^Δ^*^BT4295::BT4295^* (WT) *or B. theta*^Δ^*^tdk^*^Δ^*^BT4295::T548s^* (T548S) *in vivo*.

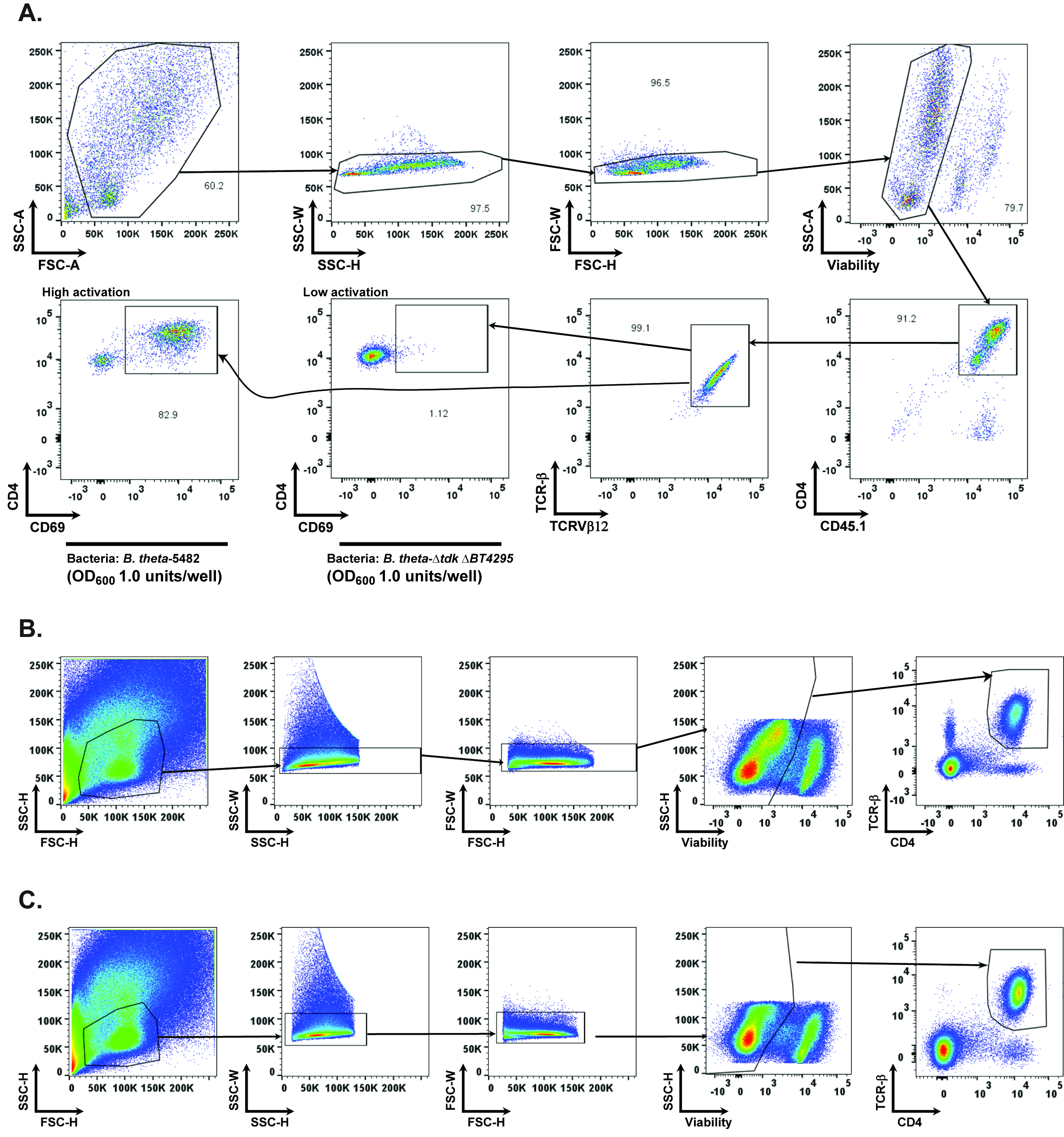
